# Ion mechanism and parameter analysis of anodal-first waveforms for selective stimulation of C-fiber nerves

**DOI:** 10.1101/2020.12.13.422609

**Authors:** Siyu He, Kornkanok Tripanpitak, Yu Yoshida, Shozo Takamatsu, Shao Ying Huang, Wenwei Yu

**Affiliations:** Graduate School of Science and Engineering, Chiba University, Japan; Omron Healthcare Co., Ltd., Japan; Engineering Product Development, Singapore University of Technology and Design, Singapore; Center for Frontier Medical Engineering, Chiba University, Japan

## Abstract

Few investigations have been conducted on selective stimulation of small-radius unmyelinated C nerves (*C*), which is critical to both recoveries of damaged nerves and pain suppressions. The purpose of this study is to understand how an anodal pulse in an anodal-first stimulation could improve *C*-selectivity over myelinated nociceptive Aδ nerves (*Aδ*), and further clarify the landscape of the solution space. The Hodgkin-Huxley (HH) model and the McIntyre-Richardson-Grill (MRG) model were used for modelling *C* and *Aδ,* respectively to analyze the underlying ion dynamics and the influence of relevant stimulation waveforms, including monopolar, polarity-symmetric, and asymmetric pulses. Results showed that polarity-asymmetric waveforms with preceding anodal stimulations benefit *C*-selectivity most, underlain by the decrease of the potassium ion current of *C*. The optimal parameters for *C*-selectivity have been identified in the low frequency band, which will benefit remarkably the designs of nociceptive nerve selective stimulation.

## Introduction

Electrical stimulations have been shown to be effective to restore functions of sensory and motor nerves [Eisenberg 1987, Sisken 1993, Ahlborn 2007, 1–3]. Especially among them, the electrical stimulations of the unmyelinated C-fiber nerves (*C*) contributes to the recovery of damaged nerves and pain suppression. As reported in [Paquette 2019, 4], tibial nerve stimulation (TNS) that activated *C* could cause statistically significant inhibiting effects on bladder overactivity, though selective stimulation of *C* has not been addressed, since not only *C*, but also Aδ-fiber nerves (*Aδ*) and Aβ-fiber nerves *(Aβ)* were activated. Stimulating the unmyelinated nerves also helps to suppress chronic pain, which is one of the main reasons for reducing the quality of life (QOL) of patients. However, there have been few effective treatments for chronic pain to date [Apkarian 2011,5].

For the selective stimulation of peripheral unmyelinated nerves, drug control [Breivik, Harald 2006, 6], spot-oriented needle stimulation or electroacupuncture [Chen, Xiao-Hong 1992, 7], laser [Plaghki, Léon 2003, 8] heat simulation [Churyukanov, Maxim 2012, 9], and the low-frequency electrical stimulation (0-20Hz) [Harris, G. W. 1969, 10] have been studied and some of them have led to clinical or medical research use. For instance, it is possible to use drugs to inactivate certain ion channels, thus select specific nerve fibers [Grill, Warren M. 1991, 11]. However, the influence and harm of drugs on human body due to addiction are also obvious [Martell, Bridget A 2007, 12]. For the spot-oriented stimulation spatial selection, in clinical practice, needles or needle electrodes (i.e. acupuncture electrodes) are inserted to specific spots that have higher spatial distribution of the *C* [Otsuru 2009, 13]. Since using acupuncture electrodes for stimulation requires professional skills, it is difficult to promote to the public use for chronic pain relief [Siyu He 2020, 14]. The laser and heat stimulation have been widely used in the laboratory-based *C* stimulation for pain-related clinical studies [Plaghki, Léon 2003, Churyukanov, Maxim 2012, 8–9]. However, same as the electroacupuncture, special devices and skills are necessary for an operation. Consequently, except the electrical stimulation, all the others are either for clinical use only, or requiring professional level skills.

Generally, when electrically stimulated, thicker *Aδ* are more likely to be excited than thin *C* [Grill 1997, 15]. The reason is that a thicker nerve has a larger membrane area, which causes a drop in electrical resistance. Moreover, myelin sheaths result in a high concentration of sodium channels between them (i.e. nodes of Ranvier), which eases the excitation of *Aδ*. Compared with *Aδ*, *C* is thin and has no myelin sheath. Both factors make it difficult to stimulate the thin *C* before the thicker *Aδ* are activated.

Nevertheless, distinct properties of the ion channels of different nerve fibers keep open the possibility of selective stimulation of *C*. There have been two categories of approaches for selective stimulations of nociceptive nerve fibers. The first one is nerve-specific frequency stimulation, which makes use of distinctive response of nociceptive nerves to different frequency bands. One commercialized device, Neurometer® (Neurotron, Baltimore, MD, USA), employs sinusoidal waves at frequencies of 5 Hz, 250 Hz, 2000 Hz to selectively stimulate *C*, *Aδ*, and *Aβ,* respectively [Pitei 1994, 16]. It demonstrates the possibility of selective stimulation by surface electrodes. However, as was reported, those low-frequency sine waves cannot selectively stimulate the *C* securely [Koga 2005, Dufour 2011,17-18]. The second category of approach is pre-pulse stimulation, which applies a preliminary stimulation to alter the states of the neuromembrane and ion channel gate variables, thus change the activation threshold of a certain type of nerves responding to a following stimulation. Bostock et al. proposed a method called QTRAC© (Institute of Neurology, London) to diagnose neuropathy according to the neuronal excitability of myelinated nerves by cathodal pre-pulse stimulation [Bostock 1998, 19]. This method shows that a preceding cathodal pulse before a main cathodal stimulation can promote the reduction of the stimulation intensity for excitation, which is an increase of the excitability of a nerve. In 1995, Grill et al. proposed that a preceding square stimulation can change the neural excitation threshold of myelinated nerves, i.e., a cathodal pre-stimulation can suppress, and an anodal pre-stimulation can promote nerve excitability [Grill 1995, 20]. However, the findings of the two studies on myelinated nerves are opposite. The contradiction might be caused by the different characteristics of their ion channel models. Along this consideration, the unmyelinated fibers might show difference in response to a preceding stimulation, which has not been addressed so far in the literature, thus needs to be investigated systematically, while compared with the *Aδ*.

On the other hand, sensory nerve action potentials (SNAPs) can show two separate deflections, i.e., double peak potentials, responding to nerve stimulation. Moreover, it has been discovered in animal experiments that only an anodal stimulation can generate action potentials in tissues such as myocardium, which was named as anode break [Ranjan 1998, 21]. It was further investigated with an axon-level theoretical framework, showing that an inward-rectifier potassium ion current is essential for an anode break [Ranjan 1998, 21]. Both membrane potential and stimulation duration affect the development of an anode break. When the membrane potential is in a more depolarized position, the potassium ion current decreases, and a current flowing into the axon is generated only at the end of an anodal stimulation to form an action potential.

Our hypothesis is that, the mechanism of anode break might work to further lower the threshold of *C*, too. In our previous study, we clarified that a low-frequency bipolar square waveform could improve the *C*-selectivity [Siyu He, 2020, 14]. However, neither the mechanism of the preceding anodal stimulation for the *C*-selectivity, nor the important parameters of the bipolar square wave have been explored.

The purpose of this study is to investigate and understand the efficacy of preceding anodal stimulation on selectively stimulating *C* and the ion channel mechanism behind. It is aimed to identify the waveform parameters that enable the selectivity of nociceptive *C* over myelinated nociceptive *Aδ* using computation models of the nerves, and further clarify the landscape of the solution space. Relevant stimulation waveforms, including monopolar, polarity-symmetric preceding cathodal bipolar, polarity-symmetric preceding anodal bipolar waveforms, and their asymmetric correspondents, were investigated with not only their behavior in ion channel gate variable – membrane potential phase portrait (which associates the nerve activation with the ion channel mechanism), but also the ratio of the excitation threshold of *C* to that of *Aδ*, defined as *Rth.* The Hodgkin–Huxley (HH) model and McIntyre-Richardson-Grill (MRG) model were employed to represent the *C* [Hodgkin 1952, 22] and the *Aδ,* respectively [McIntyre 2002, 23].

The HH model, as a classic model of neural behavior, produces predictable action potentials and reveals a wealth of information about neural signaling. It has been applied to and validated with the experiment data of squid giant axons [Best 1979, 24], and mammalian axons [Krouchev 2015, 25], and used to study the unmyelinated nerves of various organisms nowadays [Stein 1971, Schoenbach 1997, Pinto 2000, 26–28]. Regarding the MRG model, many simulations and animal experiments have demonstrated its validity as the model of motor and myelinated sensory nerves [Bhadra 2007, Bourbeau 2011, Gaines 2018, 29–31].

The rest of the paper is organized as follows. Section 2 introduces models that simulate *C* and *Aδ,* describing the time constants of each model, dynamical behavior of ion channel variables responding to different stimulation waveforms. Section 3 presents experimental results of phase portrait analysis, and exploration of the solution space for *C* selective stimulation. Section 4 gave a discussion on experiment results and models, contributions of this study, and future directions. The paper is concluded with Section 5.

## Model & Method

### 2.1 HH model and MRG model

The HH model which was first published in 1952 has changed the understanding of neuronal function [Hodgkin 1952, 22] and provides a quantitative description of action potential generation and variation, as well as the structure and function of ion channels [Catterall 2012, 32]. It reproduces the responses to electrical nerve stimulation by using differential equations with appropriate coefficients obtained in animal experiments [Hodgkin 1952, 22]. A model of *C* was proposed based on the original HH model, which could express potential propagation between nodes, thus enable the model identification even with experiment data regarding nerve conduction velocity from earlier studies of patients and animals [Bekkouche 2012, 33]. In our study, HH model, as expressed in Equation (1)–(3), was used to realize the human *C*, by tuning model parameters, such as resistivity, fiber diameter, etc. with the experiment results provided by [Tarotin 2018, Tai 2005, 34–35].

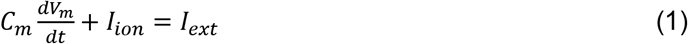

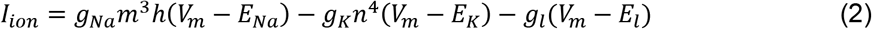

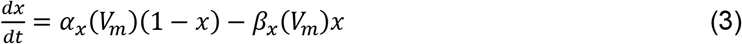

Equation (1) shows the change of transmembrane current by an external stimulation. *C_m_* is the membrane capacitance, *V_m_* is the intracellular potential, *t* is time, *I_ion_* is the ionic transmembrane current, and *I_ext_* is the externally applied stimulation current. The details of ionic transmembrane current *I_ion_* are shown in Equation (2). The ion channel gate variables *m, h* and *n* define the open status of those ion channel. *m* and *h* define the opening and closing of sodium ion channels respectively, and *n* defines the opening of potassium ion channels. *g_Na_, g_K_* and *g_l_* represent the maximal conductance of each ion channel. *E_Na_, E_k_* and *E_l_* represent the resting potential of each channel. Equation (3) shows the first order derivative of each gate variable (*x* = [*m, h, n*]), defining the open probability of each ion channel. *α* and *β* are the rate constants dependent on the transmembrane voltage, which describe the transient rates of channel gates opening and closing. The values of the parameters were shown in Appendix I. The equivalent circuit of the HH model is shown in Figure 1A.

**Figure 1.**
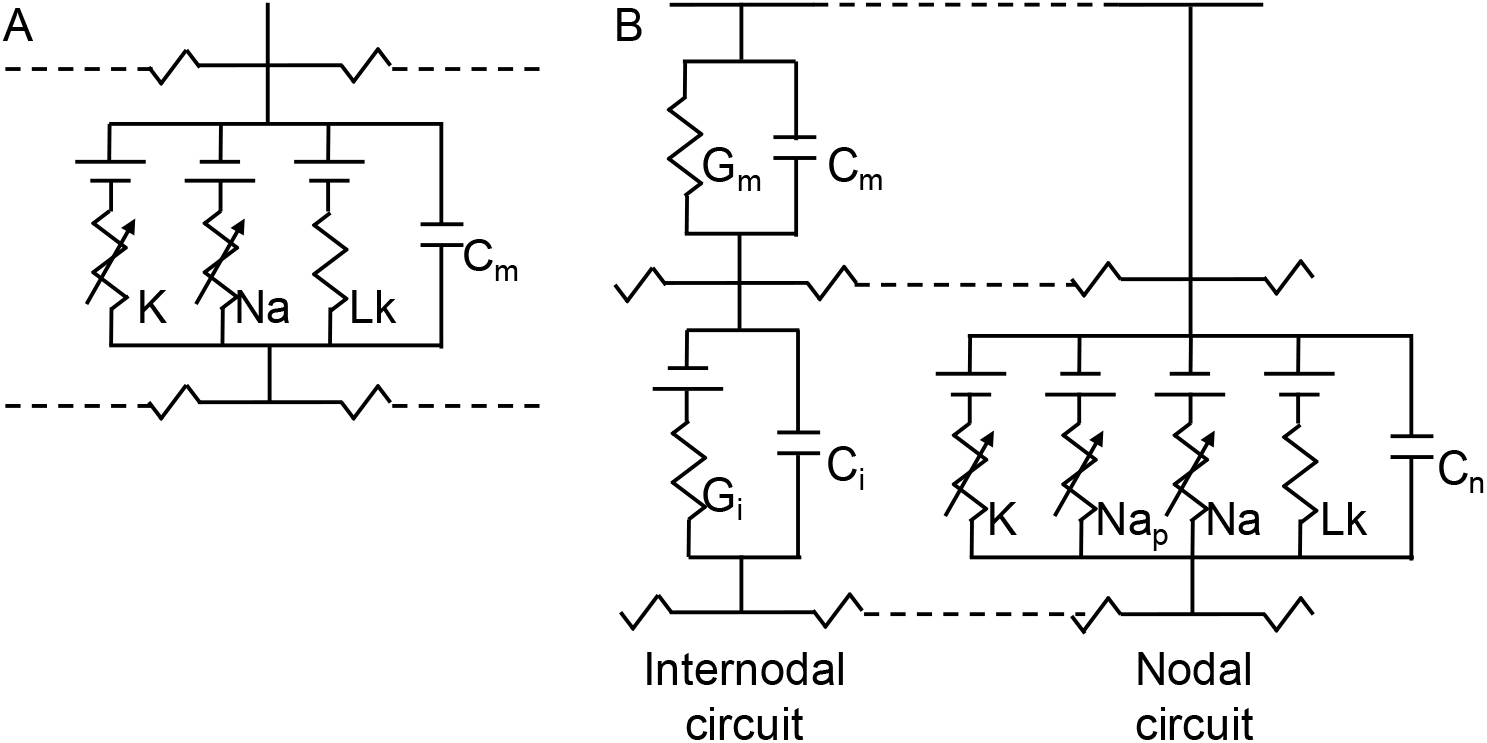
The HH (A) and MRG (B) double-cable axon model. (A) A segment of unmyelinated fibers contains sodium *(Na),* potassium *(K)* channels, and leakage resistance *(Lk)* with membrane capacitance (*C_m_*). (B) A node of Ranvier segment (i.e. nodal circuit on the right) contains fast sodium (*Na*), persistent sodium *(Na_p_),* slow potassium (*K*) channels, and leakage resistance (*Lk*) with nodal capacitance (*C_n_*). An inter-nodal segment contains resistance and capacitance of myelin sheath (*G_m_* and *C_m_*), and inter-nodal double-cable structures *(G_i_* and *C_i_*) [McIntyre 2002, 23].

Regarding myelinated nerve fibers, the FH (Frankenhaeuser, Huxley) model was first established to simulate their behavior, expressed by the experimental data on myelinated nerve fibers of frog [Frankenhaeuser 1964, 36]. Since the structure and properties of ion channels at the nodes of Ranvier in mammals are distinctly different from those of amphibians, the neural model about mammals has been further developed [Sweeney 1987, 37]. McIntyre et al. developed geometrically and electrically accurate models of mammalian motor nerve fibers, MRG (McIntyre, Richardson, and Grill) models, to study the biophysical mechanisms of axonal excitability changes and the recovery cycle regulation [McIntyre 2002, 23]. The MRG model can reproduce the experimental data about the excitation properties of mammals myelinated nerve fibers.

The MRG model, which represents *Aδ*, can be expressed using Equation (1), too, since it stems from the classical HH model. However, for different ion channels, the ionic transmembrane current *I_ion_* was expressed by the following differential equations.

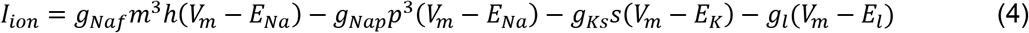

Different from equation (2), in equation (4), *Naf* means the nodal membrane fast sodium channel, *Nap* means the persistent sodium channel, *Ks* is the slow potassium channel, and *l* means the linear leakage conductance. Variable *m* and *h* are the same with those of the HH model. Variable *p* defines the opening of sodium persistent ion channels, and *s* defines the opening of slow potassium ion channels. The equivalent circuit of MRG model is shown in Figure 1B.

The key to improve the *C*-selectivity over *Aδ* is the difference in their ion channels. One of the important properties is the time constant of the ion channels. Figure 2A and B show the change of the time constant of each variable as the membrane potential *V_m_* increases, in the HH and MRG models, respectively. For the HH model, as shown in Figure 2A, the time constant of variable *m* is considerably small compared with *h* and *n,* which means the *m* exerts the greatest effect on system behavior. That is, as the membrane potential increases, even if the sodium ion channel is ready to close, as suggested by the fast-reducing *m*, the opening variable of sodium ion channel, but if *h*, the closing variable, has not reached its the maximum value, the sodium ion channel is keeping open. Prolonging this time lag means the extension of the opening time of the sodium ion channels, which has a great promoting effect on nerve fiber excitation. This is also consistent with the description in [Grill 1995, 20]. On the other hand, different from the HH model, the sodium channel time constant of the MRG model, especially the *h* in Figure 2B is smaller than that of the HH model. Therefore, the fast-changing gate variable of the sodium channel of the MRG model makes its effect time much shorter than that of the HH model. This leads to the improvement of the excitability of the *C*, which may even exceed that of the *Aδ*, providing the possibility to reverse their threshold.

**Figure 2.**
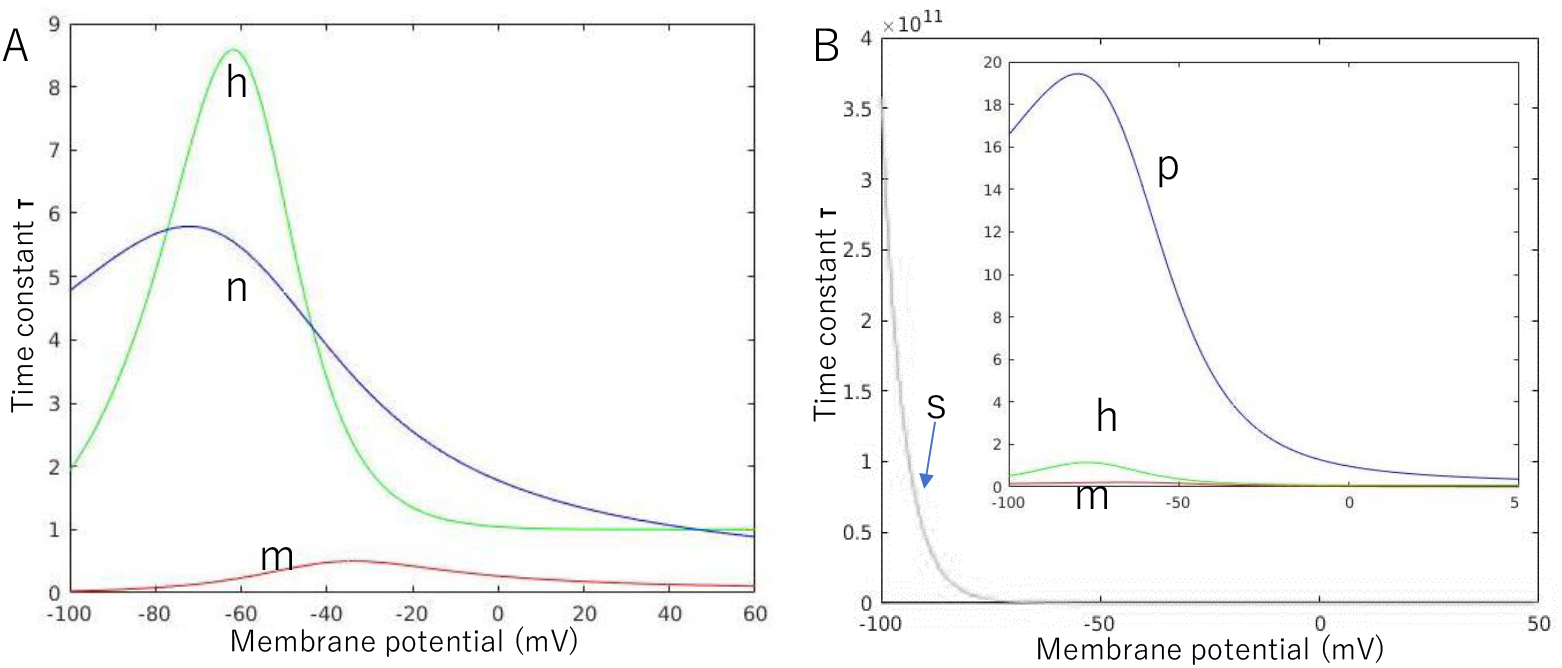
The time constants of each ion channel in HH(A) and MRG(B) model

### 2.2 Phase portrait analysis

Phase portrait could visually reveal dynamics of multi-variate data series in a phase plane. Figure 3 shows an example of phase portrait of the HH model, when no external stimulus was given. The red curve plots the isocline of *dm/dt*=0, and the blue curve plots the isocline of *dV_m_/dt*=0. The three intersections *a, b* and *c* (i.e. the points meet both *dm/dt*=0 and *dV_m_/dt*=0, at which the status does not change over time), stands for the rest state, activation threshold and the peak of action potential, respectively [Doug Dean 1983, 38]. The positional relationship of the three intersections in the phase plane directly affect the excitability of the nerve fibers. The dynamical changes of the phase portrait are subject to the time constants of each ion channel of the nerve model. In this study, the focus was put on the phase of *m* and its membrane potential *V_m_*, while ignoring other variables. The reason is that the time constants of the rest are much greater than that of *m*, which means that they can be considered as constants with respect to that of *m* within an infinitesimal time.

**Figure 3.**
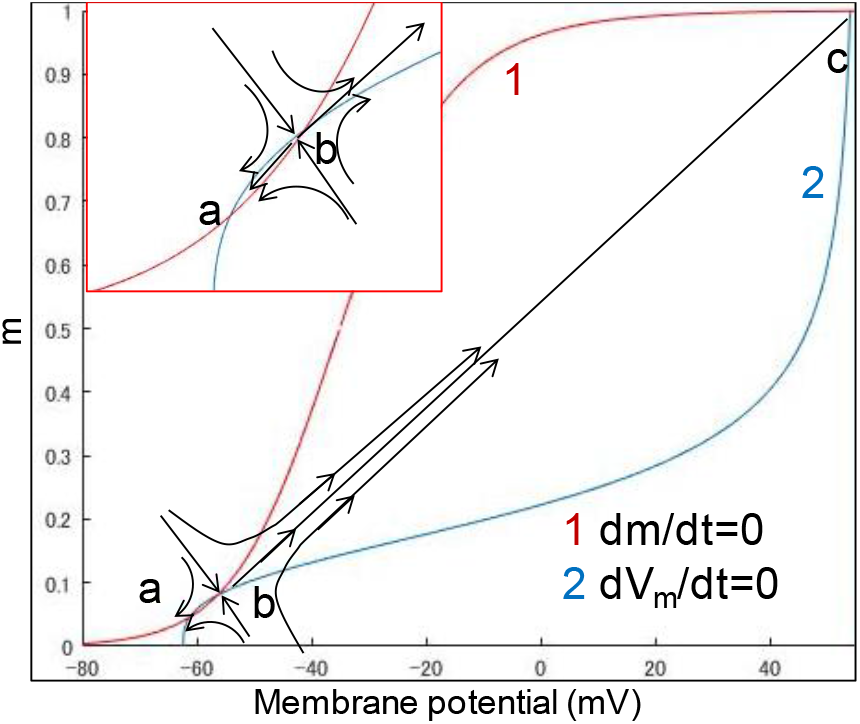
An example of phase portrait of the HH model. The figure shows the dynamic changes of HH model in rest state. The red curve *1* plots the resting state isocline of *dm/dt*=0, and the blue curve *2* plots the resting state isocline of *dV_m_/dt*=0. The three points *a, b* and *c* mean the rest point. Point *a, b* and *c* represent the situation of rest, activation threshold and peak of action potential. Arrows denote the trends of state transition. Since point *b* is a saddle point, the state point will move closer to point *a* before crossing the saddle point *b* and move to point *c* after crossing *b*.

### 2.3 Evaluation and analysis

To assess selective stimulation of *C*, excitation threshold of *C* (*Th_HH_*), that of MRG fiber nerves (*Th_MRG_*), and their ratio, *R_th_* (equation (5)) were used.

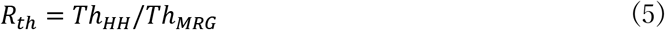

*R_th_* was used to analyze which nerve fiber model is more likely to be excited by a certain stimulation waveform. *R_th_* < 1 and *R_th_* > 1 implies that *C* is more favored than *Aδ* and *Aδ* is more favored than *C, respectively.*

### 2.4 stimulation schemes

Figure 4 shows an example of the waveform under this study. The amplitudes of the anodal and cathodal stimuli are *V_a_* and *V_c_*, respectively, whereas the durations of these two are denoted using *t_a_* and *t_c_*, respectively. In each period, there is an inter-stimulus interval (ISI) between an anodal stimulus and a cathodal one. In this paper, the following effects of the stimulation waveforms on the *C*-selectivity were investigated: 1) the effect of total duration and polar precedence (i.e., anodal-first or cathodal-first) of the waveform. 2) the influence of inter-stimulus interval (ISI) between an anodal and a cathodal stimulus. 3) the effect of polarity asymmetry ratio (PAR, e.g., PAR 1:9 means the duration of cathodal stimulus is 9 times of that of anodal stimulus, in the case of anodal-first stimulation). The anodal stimulus increases the concentration of cations (+) of tissues under the anode, and the cathodal stimulus increases the concentration of anions (-) of tissues under the cathode. Only the waveforms with charge balanced anodal stimulus and cathodal stimulus were investigated in this study.

**Figure 4.**
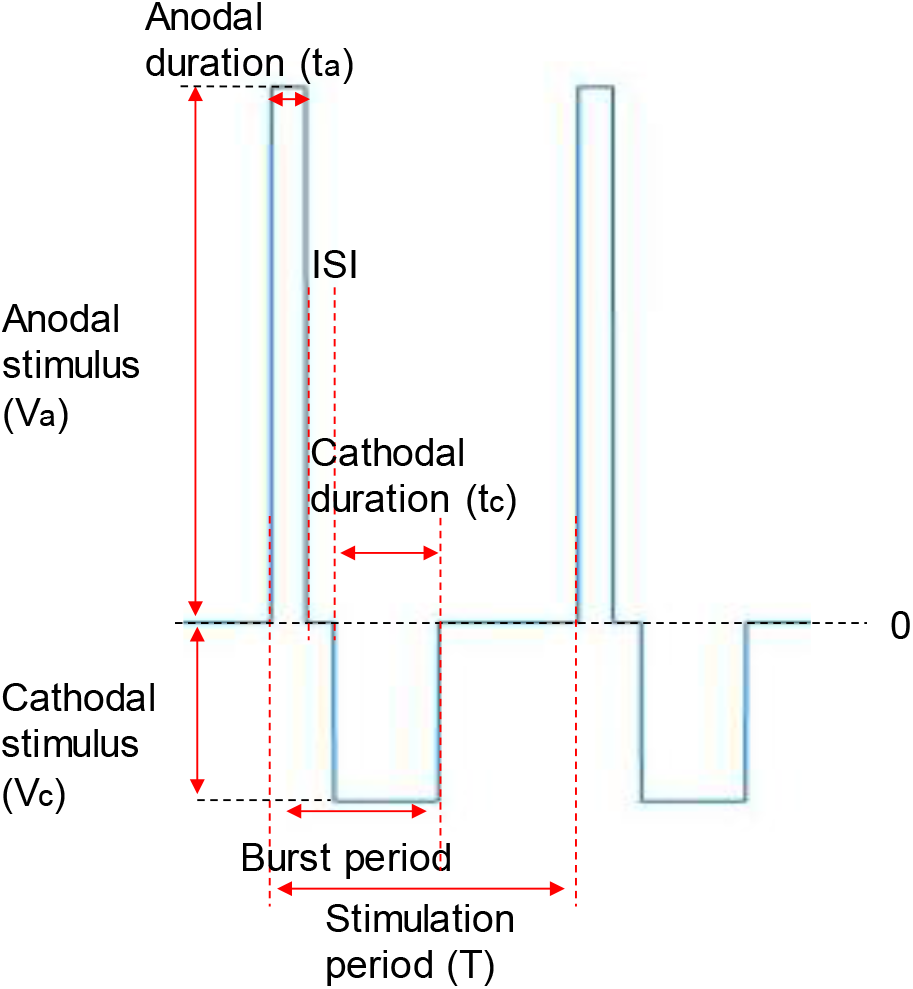
An example of the stimulation waveform. Each period contains an anodal stimulus and a cathodal stimulus. The polarity asymmetry ratio (PAR) is the quotient of the duration of the preceding stimulus and that of the following stimulus. The inter-stimulus interval (ISI) describes the no stimulus duration between the preceding and following stimulus.

### 2.5 Simulation experiments

Two categories of experiments with the simulation models were conducted: a) experiments to explore underlying ion mechanism of improving *C*-selectivity by the anodal-first stimulation, in which, focus was put on not only behavior of the *m* and *V_m_*, but also *n, h* and ion current in the phase plane; b) experiments to explore optimal stimulation parameters, in which the polar-precedence, duration, ISI, PAR and frequency were tested.

Both the HH model and MRG model used in this study contains 21 nodes. The distance between each two nodes is 10 μm and 220 μm for the HH and MRG model, respectively. Stimuli were given to a point of the 10^th^ node, at intensity of current density. Both models were implemented in Matlab 2013a, with the 4th-order Runge–Kutta (RK4) as the numerical method to solve the differential equations of the HH and MRG models. The unmyelinated and myelinated nerve models were validated with animal experimental data [Siyu He, 2020, 14].

## Results

### 3.1 Changes of activation threshold in phase portrait

Figure 5 shows the phase portrait of the HH models, for the sodium ion channel variable *m* and the membrane potential *V_m_*, (A), in their rest state (*h*: 0.59790, *n*: 0.31708), and (B), (C), (D), after anodal stimulation with duration of 2 ms, 6 ms and 10 ms, respectively, all at stimulation strength 11 μA/cm^2^. The excitability of nerves can be identified in the phase plane too. When a saddle point *b* is existing, as shown in Figure 5B, a smaller distance between two intersections, *Δd_int (a* 2-dimension vector *(ΔV_m_, Δm)* containing difference in intracellular potential *V_m_*, and difference in gate variable *m*) denotes an easier excitation of the nerve. On the other hand, as shown in Figure 5C-D, there is no intersection between the two isocline curves. In this case, the shortest distance between the two separate isocline curves, *Δd_sep* might reflect the excitation of the nerve. For the situations shown in Figure 5, (A) *Δd_int* =(2.7, 0.01402), (B) *Δd_int* =(2.0, 0.00924), (C) *Δd_sep* =(0.1,0.00057), (D) *Δd_sep* =(0.1,0.00089).

**Figure 5.**
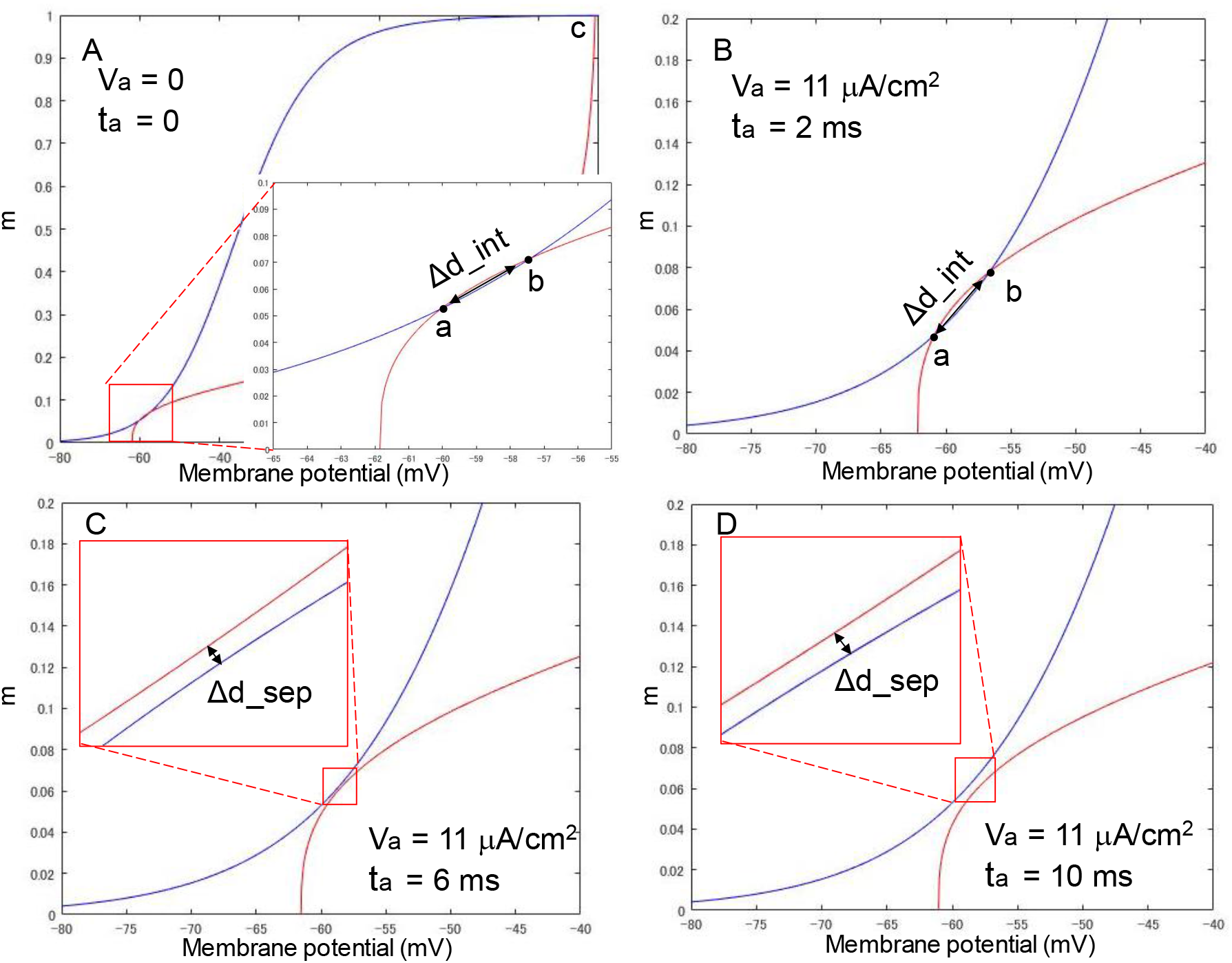
Phase portrait of the HH model with anodal stimulations of different durations (V_a_ = 11 μA/cm). (A) phase portrait of the rest status, (B) t_a_ = 2 ms. (C) t_a_ = 6 ms. (D) t_a_ = 10 ms.

### 3.2 The effect of anodal-first anodal stimulation

#### 3.2.1 Comparing the effect of anodal-first stimulation on the HH and MRG in phase plane

Figure 6 shows the phase portrait of the HH model and MRG model with different stimulation waveforms, which are illustrated by A1-F1: (A1) no stimulus, (B1) a cathodal-only stimulus with strength 162 μA/cm^2^ and duration 10 ms, (C1) a stimulus with PAR 1:1, strength of cathodal stimulus 86 μA/cm^2^ and total duration (duration of anodal and cathodal stimulus) 20 ms, (D1) a stimulus with PAR 1:9, strength of cathodal stimulus 20 μA/cm^2^ and total duration 100 ms, (E1) a stimulus with PAR 9:1, strength of cathodal stimulus 148 μA/cm^2^ and total duration 100ms, and (F1) a pre-pulse stimulus with strength 75 μA/cm^2^ and duration 90 ms, and a cathodal stimulus with strength 225 μA/cm^2^ and duration 10 ms. Note, for (F1), the intensity of preceding cathodal stimulus and following cathodal stimulus was set up according to the experiment of Bostock [Bostock 1998, 19]. For the other waveforms, the intensity was set up as their threshold strength, at which the node could reach its threshold. The phase portraits of the HH model with the stimulation waveforms are shown in Figure 6A2-F2, and those of the MRG model are shown in Figure 6A3-F3. Same with Figure 5, while rest point *a* and saddle point *b* maintain as the intersections of two isocline curves of the MRG model (Figure 6A3-F3), they disappear in phase portraits of the HH model (Figure 6C2-D2). Following the definition of distance in section 3.1, *Δd_int* and *Δd_sep* were used to analyze the responses of the HH model and MRG model, to different stimuli.

**Figure 6.**
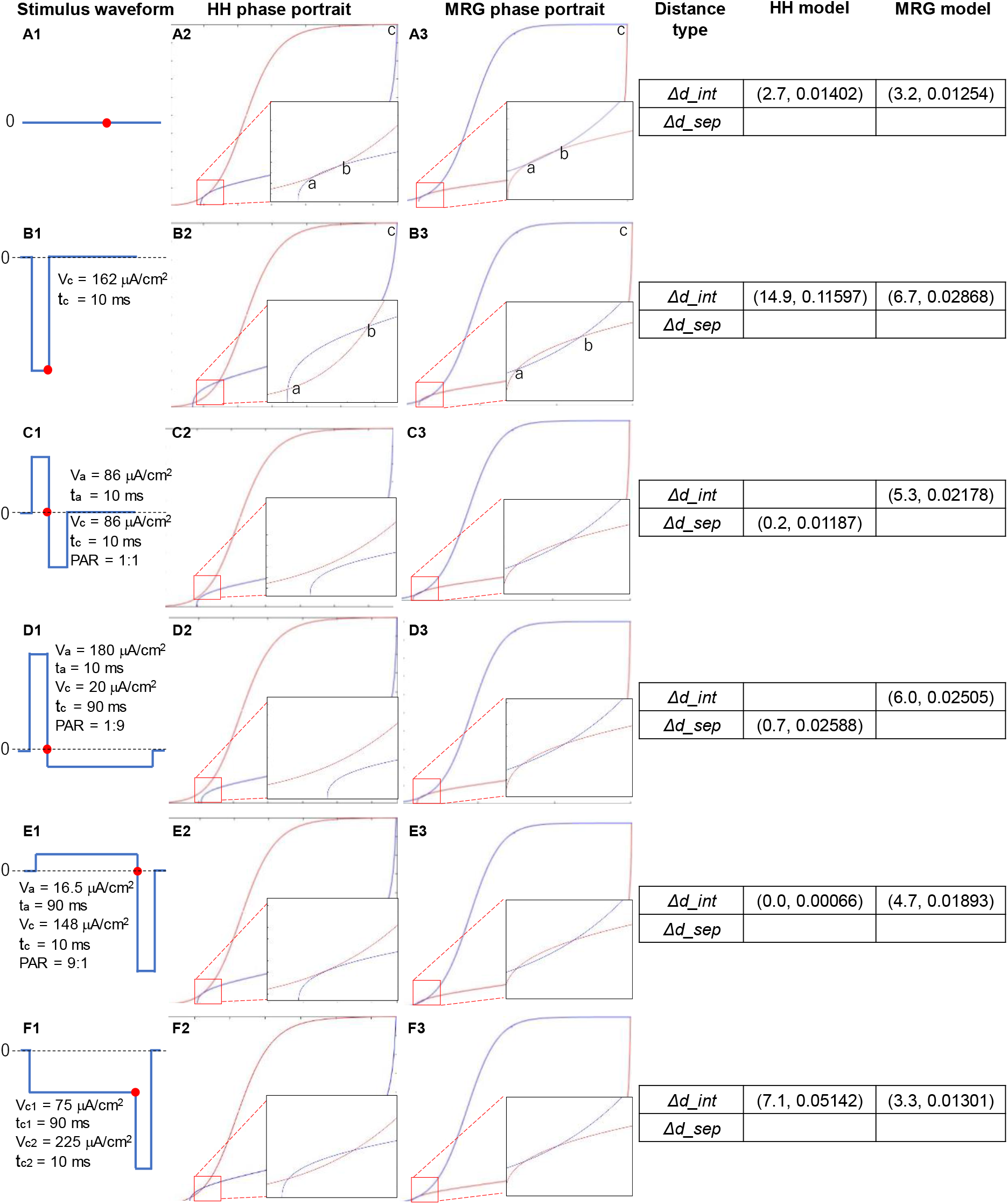
Phase portraits of dynamical behavior of the HH model and MRG model responding to different stimuli. (A1)—(F1) illustration of stimulation waves: (A1) no stimulus, (B1) a cathodal-only stimulus with strength 162 μA/cm^2^ and duration 10ms. (C1) a stimulus with PAR 1:1, strength of cathodal stimulus 86 μA/cm^2^ and total duration 20 ms. (D1) a stimulus with PAR 1:9, strength of cathodal stimulus 20 μA/cm^2^ and total duration 100 ms. (E1) a stimulus with PAR 9:1, strength of cathodal stimulus 148 μA/cm^2^ and total duration 100 ms. (F1) a pre-pulse stimulus with strength 75 μA/cm^2^ and duration 90 ms, and a cathodal stimulus with strength 225 μA/cm^2^ and duration 10 ms. (A2)—(F2) the phase portraits of the HH model. (A3)—(F3) the phase portraits of the MRG model. The 4^th^ column: *Δd_int* and *Δd_sep* as a result of different stimulation

#### 3.2.2 Changes of Ion channel variables and ion current caused by the preceding anodal stimulation

To uncover the mechanism of anodal-first bipolar simulation, membrane potential and the current of ion channels were further investigated. Figure 7 shows the changes of (A) total transmembrane ion current, (B) ion current in sodium (Na) channel, and (C) ion current in potassium (K) channel, with regard to membrane potential, responding to different preceding anodal stimulations but the same following cathodal stimulus. The preceding anodal stimulation has the same parameters as those described in (A1), (C1), (D1), and (E1) of Figure 6, i.e. no stimulus, stimuli with PAR 1:1, 1:9 and 9:1, respectively, while the cathodal stimulus is identical with that described in Figure 6B1.

A red box in each graph of Figure 7 shows the change of ion current before reaching its threshold (prethreshold phase). The black dots denote where the membrane potential exceeds the threshold, *ΔI* shows the difference between the ion current of each polarity asymmetric stimulation waveform and that of no anodal stimulation at its threshold strength. The results of each *ΔI* are 36.94 μA, 24.53 μA, 18.24 μA, 9.35 μA, and 5.18 μA, corresponding to the simulation with PAR 1:9, 1:1,2:1,7:1,9:1, respectively. Because, as shown in Figure 7B, there is no difference between the sodium ion currents corresponding to different stimulation waveforms (maximal *ΔI_Na_*: 4.50 μA at PAR=1:9, average *ΔI_Na_*: 2.58 μA), it is clear that, the difference in transmembrane ion current (maximal *ΔI*: 43.98 μA at PAR=1:9, average *ΔI*: 25.24 μA) is mainly due to the change of potassium ion current (maximal *ΔI_K_*: 48.48 μA at PAR=1:9, average *ΔI_K_*: 27.82 μA), as shown in Figure 7C.

**Figure 7.**
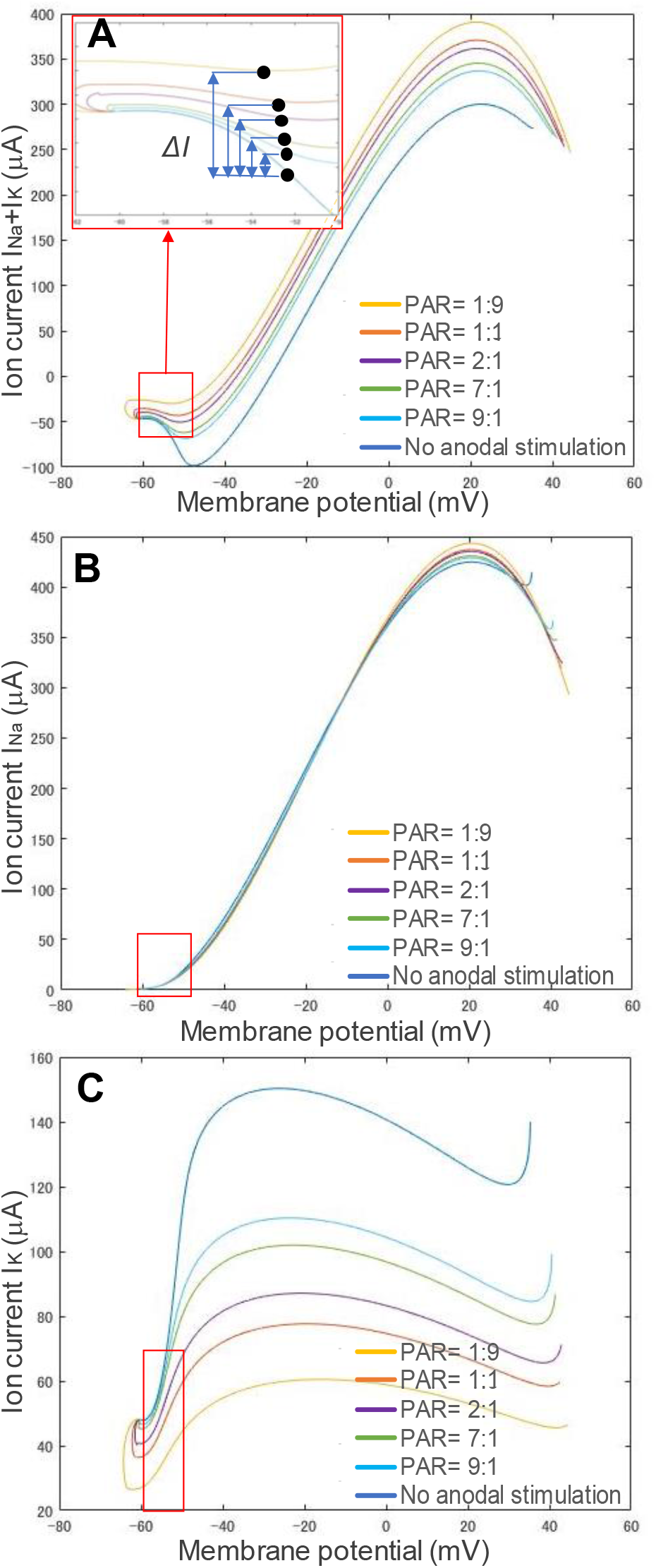
The changes of ion channel currents with respect to membrane potential responding to different preceding anodal stimulus and one identical following cathodol stimulus. (A) transmembrane ion current: *I_Na_* + *I_K_* (B) current of sodium ion channel: *I_Na_* (C) current of potassium ion channel: *I_K_*. The other parameters of the input anodal stimuli are the same as those of A1, C1, D1, and E1 in Figure 6, respectively. A red box in each graph shows the change of ion current before reaching the membrane potential threshold (pre-threshold phase).

Though, the ion channel variables *h*, and *n*, were not focused in most simulation studies, they are playing an important role in the neural dynamics. Figure 8 shows the changes of the ion channel variables (A) *m* (B) *h* (C) *n* of the HH model with respect to membrane potential, responding to different stimulation waveforms: 1) a cathodal-only stimulus with strength 162 μA/cm^2^ and duration 10 ms, 2) a stimulus with PAR 1:1, strength of cathodal stimulus 86 μA/cm^2^ and total duration 20 ms, 3) a stimulus with PAR 1:3, strength of cathodal stimulus 46 μA/cm^2^ and total duration 40ms, 4) a stimulus with PAR 1:6, strength of cathodal stimulus 27 μA/cm^2^ and total duration 70 ms, 5) a stimulus with PAR 1:9, strength of cathodal stimulus 20 μA/cm^2^ and total duration 100 ms.

**Figure 8.**
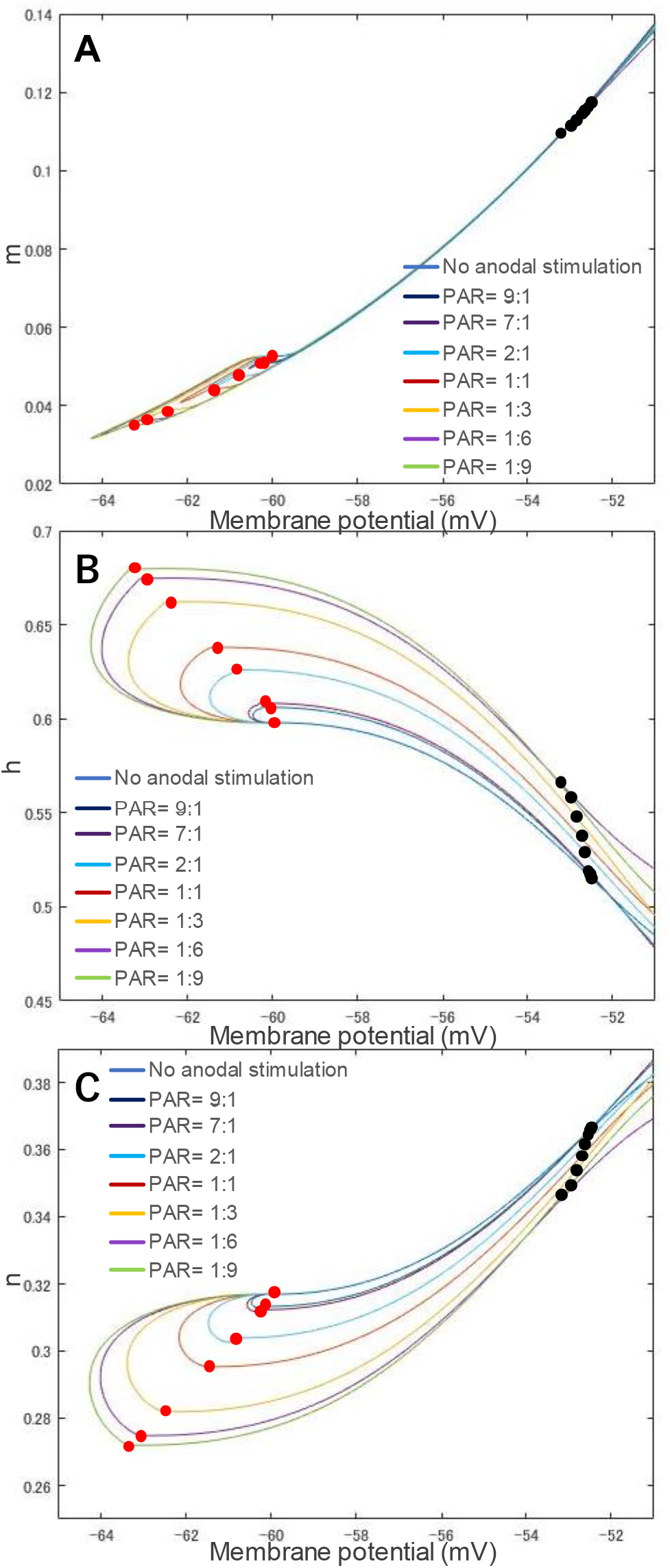
The related changes of each ion channel variable and membrane potential. (A) opening of sodium ion channel: *m.* (B) closing of sodium ion channel: *h.* (C) opening of potassium ion channel: *n*. A red dot in each curve on the left side of each figure means the beginning of cathodal stimulation, and a black dot on the right side denotes where the membrane potential exceeds its threshold. Figure 9 shows the changes of *I_Na_* in (A) and *I_K_* in (B), with respect to membrane potential calculated from the corresponding ion channel variables in Figure 8, in the pre-threshold phase. The input stimulus waveforms are the same as those in Figure 8, at their corresponding threshold strength. This is different from the experiment shown in Figure 7, in which different preceding anodal stimulus and same following cathodal stimulus were used to investigate the effect of preceding anodal stimulation to *C*-selectivity. Despite the difference in waveforms and threshold strength, there is no significant difference between the sodium (Na) ion currents (average: 10.22 μA, standard deviation: 1.15 μA), though there is a significant gap in *I_K_* (average: 72.56 μA, standard deviation: 9.25 μA), where the membrane potential exceeds its threshold. A lower PAR 1:9 results in a higher *I_Na_*, or a smaller *I_K_*. After the membrane potential exceeds the threshold potential, as shown in Figure 9C, the potassium (K) ion currents *(IK)* showed a different spatial relationship from those in pre-threshold phase, i.e., the PAR (including no anodal stimulation) does not affect to the spatial relationship of ion currents in post-threshold phase.

Note that, for each waveform, the stimulus intensity was determined at threshold strength. A red dot in each curve shows the beginning of its cathodal stimulus, and a black dot denotes where the membrane potential exceeds the threshold, namely the onset to excite. As PAR decreases, the threshold is reduced. A different preceding anodal stimulation gives its following cathodal stimulus a different initial state (denoted by a red dot). A stronger anodal stimulus (i.e., a lower PAR such as 1:9), causes a bigger deviation from its original position (i.e., the beginning position of no anodal stimulus case) in the phase portraits.

### 3.3 The influence of waveform parameters on the *C*-selectivity

The parameter setups are shown in TABLE 1 to explore optimal stimulation parameters.

**Table 1.**
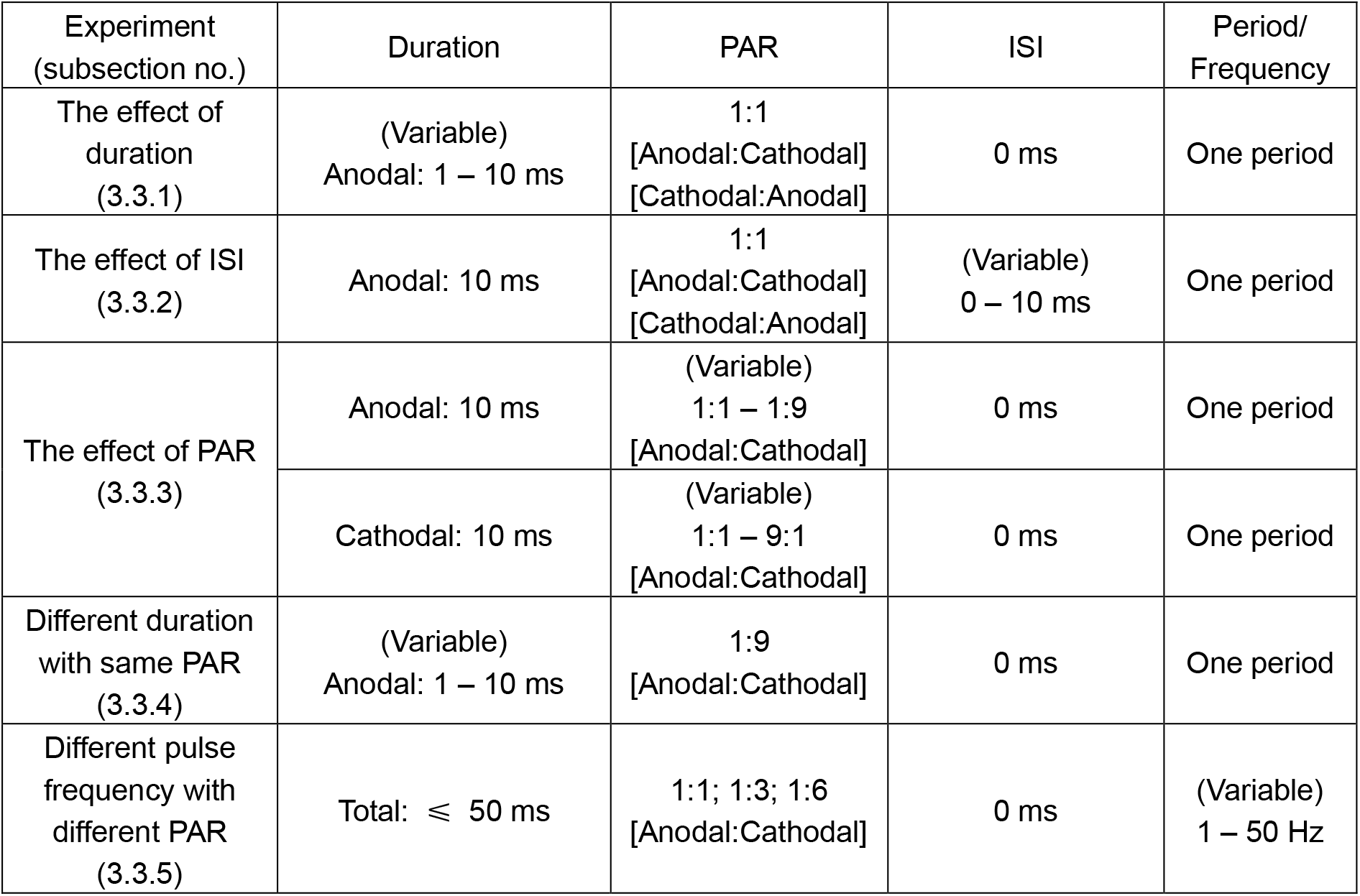
Parameter Setups For C-Selectivity Stimulation

| Experiment<br>(subsection no.) | Duration | PAR | ISI | Period/<br>Frequency |
| --- | --- | --- | --- | --- |
| The effect of<br>duration<br>(3.3.1) | (Variable)<br>Anodal: 1 – 10 ms | 1:1<br>[Anodal:Cathodal]<br>[Cathodal:Anodal] | 0 ms | One period |
| The effect of ISI<br>(3.3.2) | Anodal: 10 ms | 1:1<br>[Anodal:Cathodal]<br>[Cathodal:Anodal] | (Variable)<br>0 – 10 ms | One period |
| The effect of PAR<br>(3.3.3) | Anodal: 10 ms | (Variable)<br>1:1 – 1:9<br>[Anodal:Cathodal] | 0 ms | One period |
|  | Cathodal: 10 ms | (Variable)<br>1:1 – 9:1<br>[Anodal:Cathodal] | 0 ms | One period |
| Different duration<br>with same PAR<br>(3.3.4) | (Variable)<br>Anodal: 1 – 10 ms | 1:9<br>[Anodal:Cathodal] | 0 ms | One period |
| Different pulse<br>frequency with<br>different PAR<br>(3.3.5) | Total: ≤ 50 ms | 1:1; 1:3; 1:6<br>[Anodal:Cathodal] | 0 ms | (Variable)<br>1 – 50 Hz |

#### 3.3.1 The effect of duration

Figure 10 shows the changes of the threshold strength of the HH and MRG model, as the anodal stimulus duration changes from 1 ms to 10 ms. Bipolar symmetrical square waves were used. Polar precedence was investigated by comparing the cathodal-first (C.F.), and anodal-first (A.F.). The duration-threshold strength curve of the HH model, and that of the MRG model had an intersection at about 4 ms duration in Figure 10A (for A.F. case), i.e., the threshold strength of the two models are reversed after 4 ms. But there is no intersection in Figure 10B (for C.F. case), i.e., the threshold of the two models remain constant with long duration. In Figure 10C, the threshold ratio *(R_th_)* decreases as duration increases. The *R_th_* could be smaller than 1.0 only in the A.F stimulation case.

**Figure 9.**
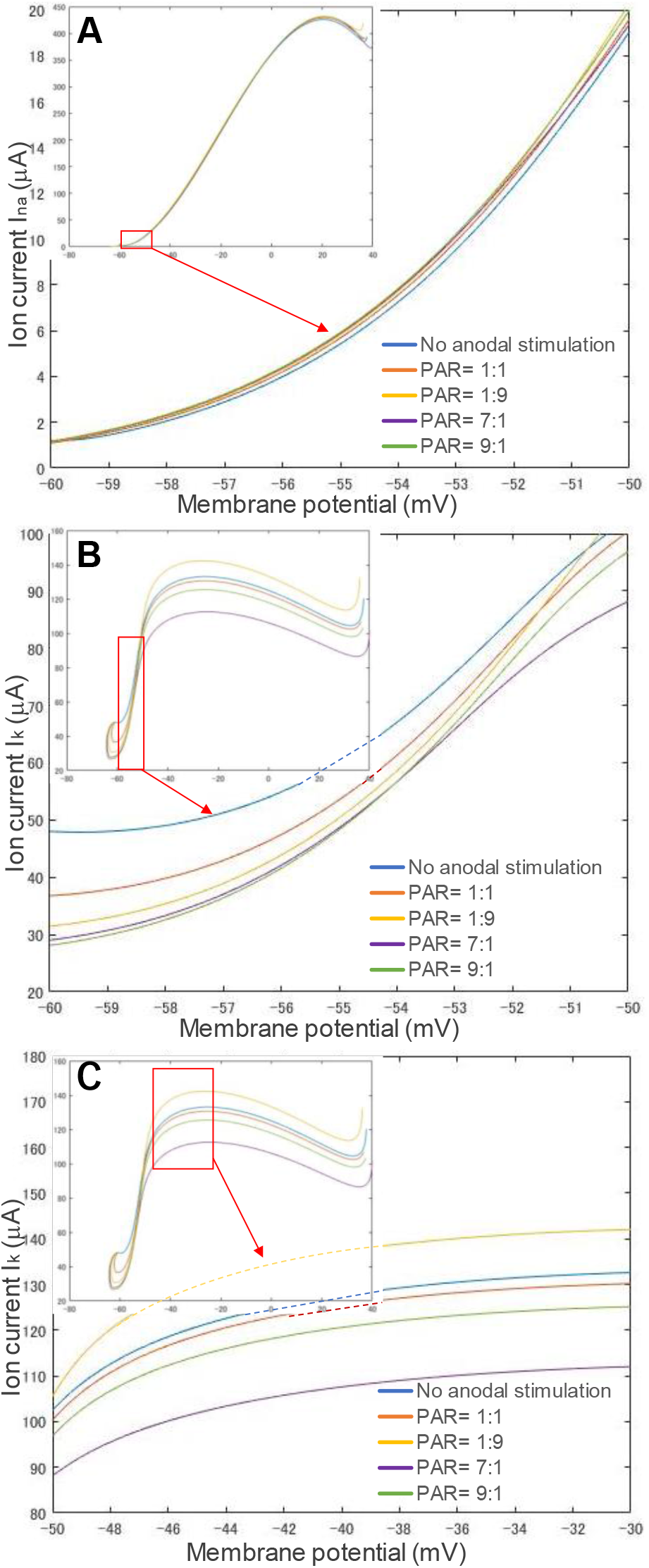
The changes of sodium *(I_Na_*) and potassium (*I_K_*) ion channel current and membrane potential responding to different stimulation waveforms. (A) sodium channel current, (B) potassium channel current before firing, (C) potassium channel after firing

**Figure 10.**
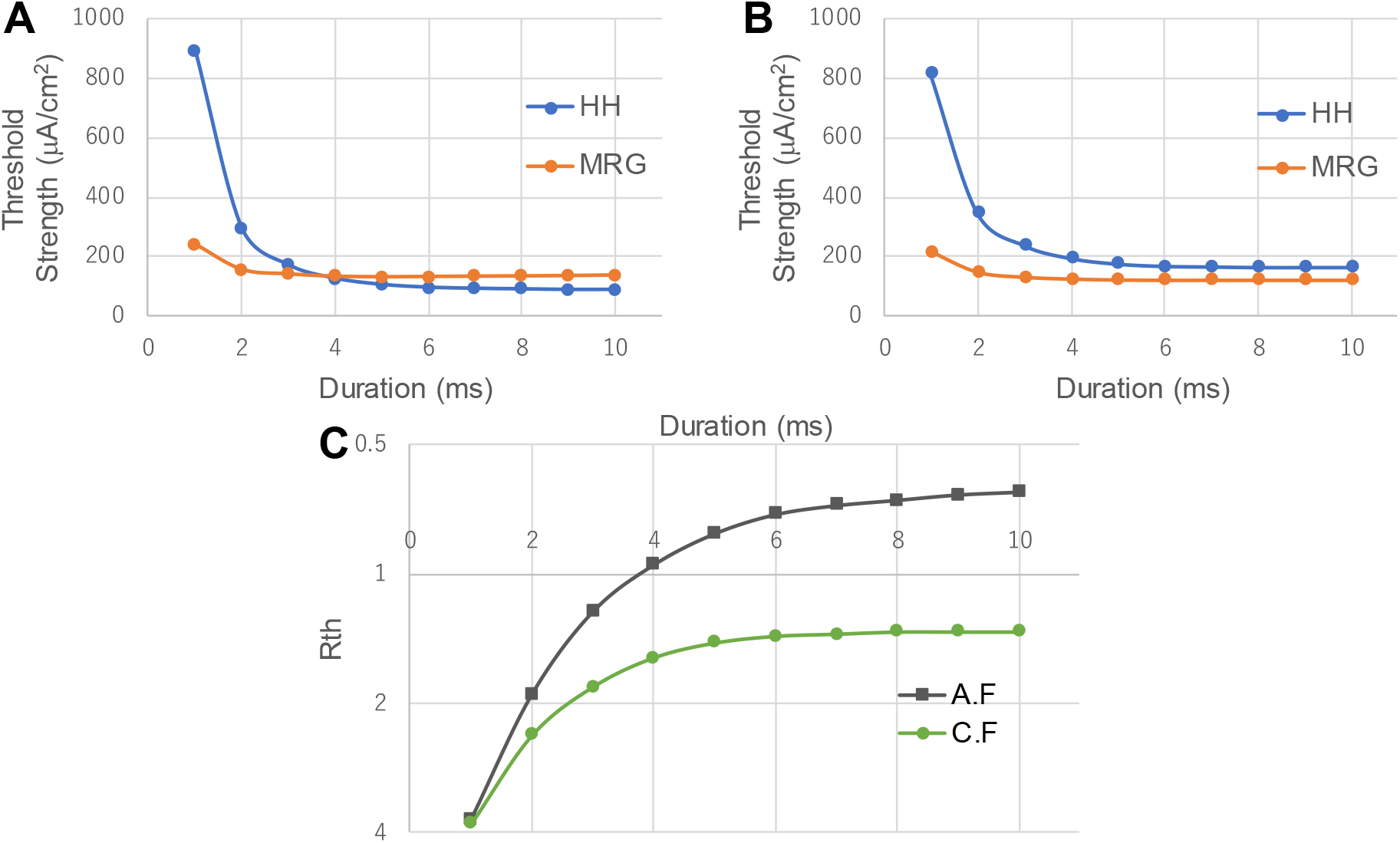
Stimulation with different durations bipolar symmetrical waveform. The horizontal axis shows anodal stimulus duration. The vertical axis represents the threshold strength. (A) anodal stimulus before cathodal stimulus (A.F: anode first). (B) cathodal stimulus before anodal stimulus (C.F: cathode first). (C) the threshold ratio between HH model and MRG model *(R_th_)* with different duration values. Fibers were stimulated with bipolar symmetrical square waves with no ISI.

#### 3.3.2 The effect of ISI

Figure 11A-B show the effect of the length of ISI between an anodal stimulus and its following cathodal stimulus on the threshold changes of HH model and MRG model. The threshold of the HH model changed significantly with ISI when in the A.F. stimulation. The threshold value starts to rise when the ISI is larger than 2 ms, and after 5 ms, it exceeds the threshold value of the MRG model. In addition, the thresholds of the HH model and the MRG model did not change significantly when the cathodal stimulation was ahead. Figure 11C shows the change in *R_th_* with ISI. Note that the threshold of the HH model when the anodal pulse is ahead will eventually be even higher than the threshold when the cathode is ahead. Figure 11D shows the phase portrait when the ISI is 4 ms A.F. The distance between two different curves *ΔdHH_sep* is (0.1,0.00154).

**Figure 11.**
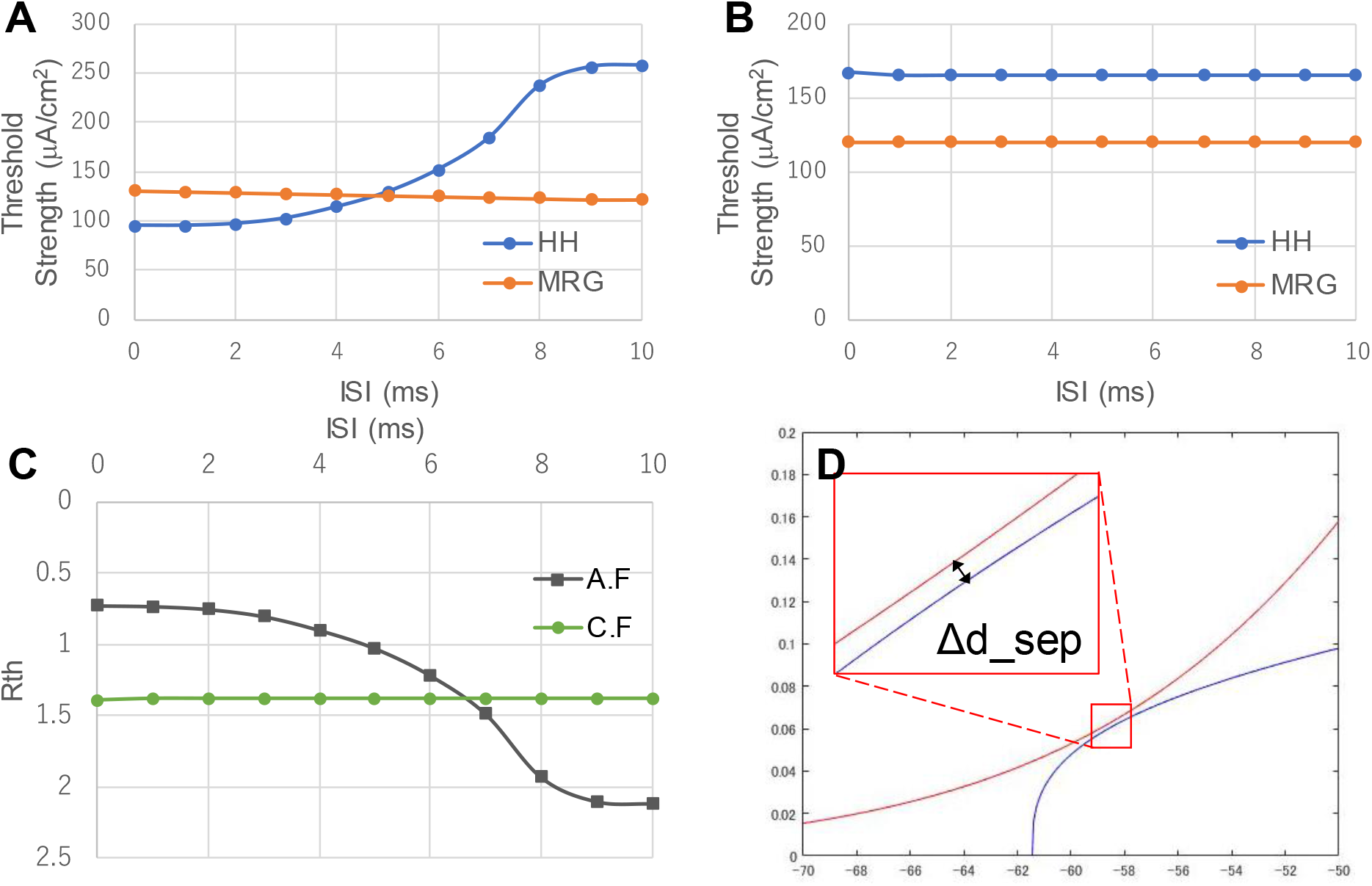
Relationship between ISI and threshold strength. (A) ISI-threshold strength relationship in A.F. stimulation case. (B) ISI-threshold strength relationship in C.F. case. (C) ISI-threshold ratio *R_th_* relationship of bother A.F., and C.F. cases. (D) the phase portrait when the ISI is 4 ms with A.F. Fibers were stimulated with bipolar symmetrical square pulses with 10 ms anodal duration.

#### 3.3.3 The effect of PAR

The threshold ratio *R_th_* changes significantly with the anodal and cathodal stimulus duration ratio, when keeping charge balance of anodal and cathodal stimuli. As an example, if anodal duration is 10 ms and cathodal duration is 90 ms, then to keep the charge balance, anodal strength needs to be 9 times of cathodal strength). Figure 12 shows the changes of threshold strength in (A) and (B), ratio *R_th_* in (C), and required charge in (D) with different PAR values. It can be seen from Figure 12A-B that the *Aδ* simulated by the MRG model has smaller changes than the *C* simulated by the HH model. In contrast, the threshold strength of *C* changes significantly. As the duration of the cathodal stimulus increases, the anodal stimulation has higher strength than the cathodal stimulus, and the stimulation threshold of the *C* gradually decreases, which suggests potential of polarity asymmetric stimulation for *C*-selectivity.

**Figure 12.**
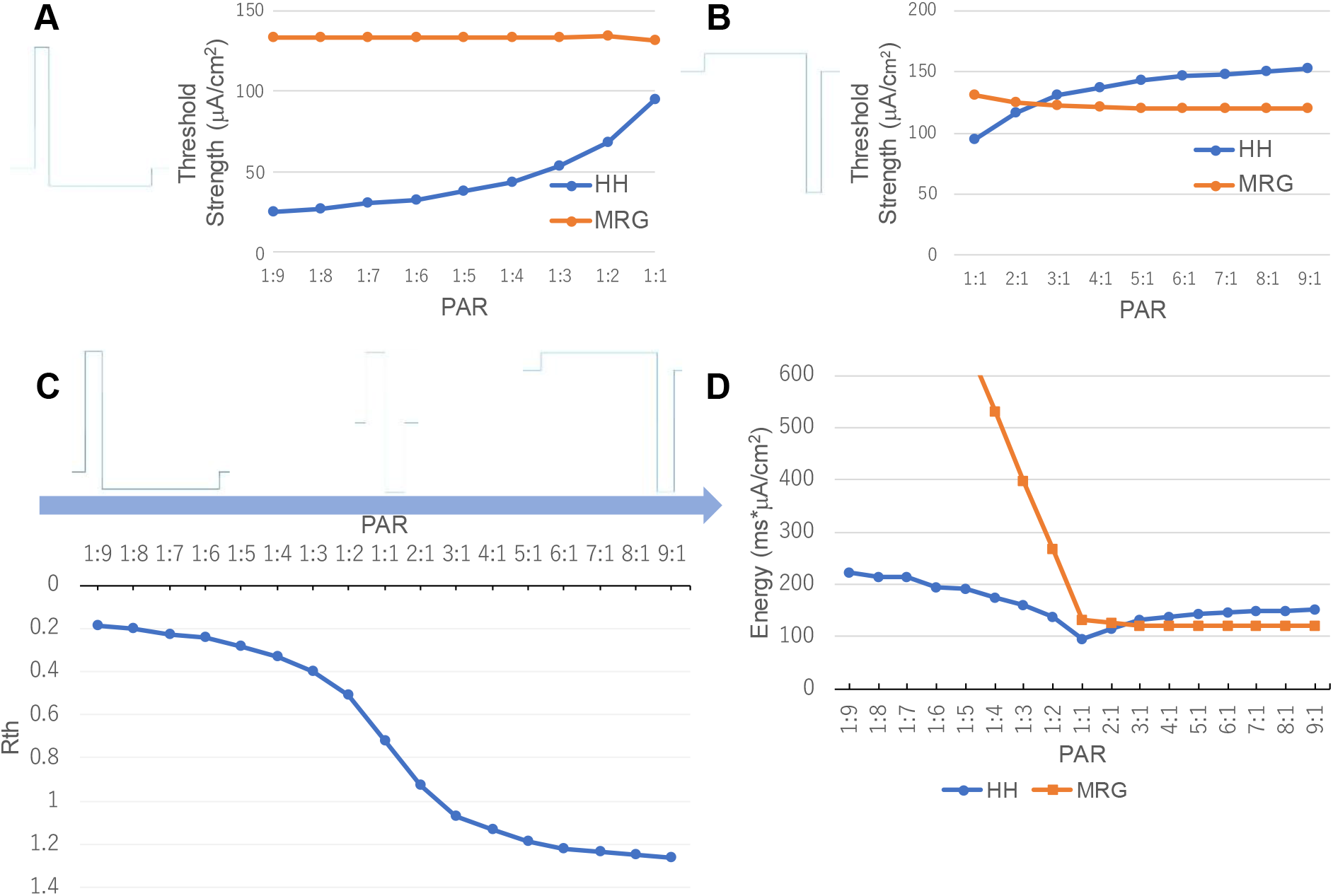
Relationship between PAR and threshold strength. (A) the strength of simulated activation threshold of *C* and *Aδ* stimulated by the waveform which the anodal stimulus is higher than the cathodal stimulus. (B) the strength of simulated activation threshold of *C* and *Aδ* stimulated by the waveform which the anodal stimulus is lower than the cathodal stimulus. (C) the change of *R_th_* with different PAR. (D) the charge of different PAR. In this experiment, the ISI between anodal and cathodal stimuli is zero.

#### 3.3.4 Different duration with same PAR

Figure 13 shows the threshold strength and threshold ratio of the HH and MRG model when stimulated with anodal stimulus with different anodal duration values at PAR 1:9. Note the total duration of the stimulus is 10 times of anodal stimulus. Only A.F. waveforms were tested, since C.F. waveforms unable to excite *C* before *Aδ*, as shown in Figure 10 in section 3.3.1. In Figure 13A, threshold strength of the MRG model does not change significantly as duration increases. In addition, the threshold strength of the HH model decreases rapidly when duration is smaller than 4 ms, then slowly decreases from 5 ms, and becomes constant from 8 ms. As shown in from Figure 13B, The *R_th_* changes from 0.62320 to 0.15833, favoring more *C*-selectivity, as anodal duration increases from 1 ms to 10 ms,

**Figure 13.**
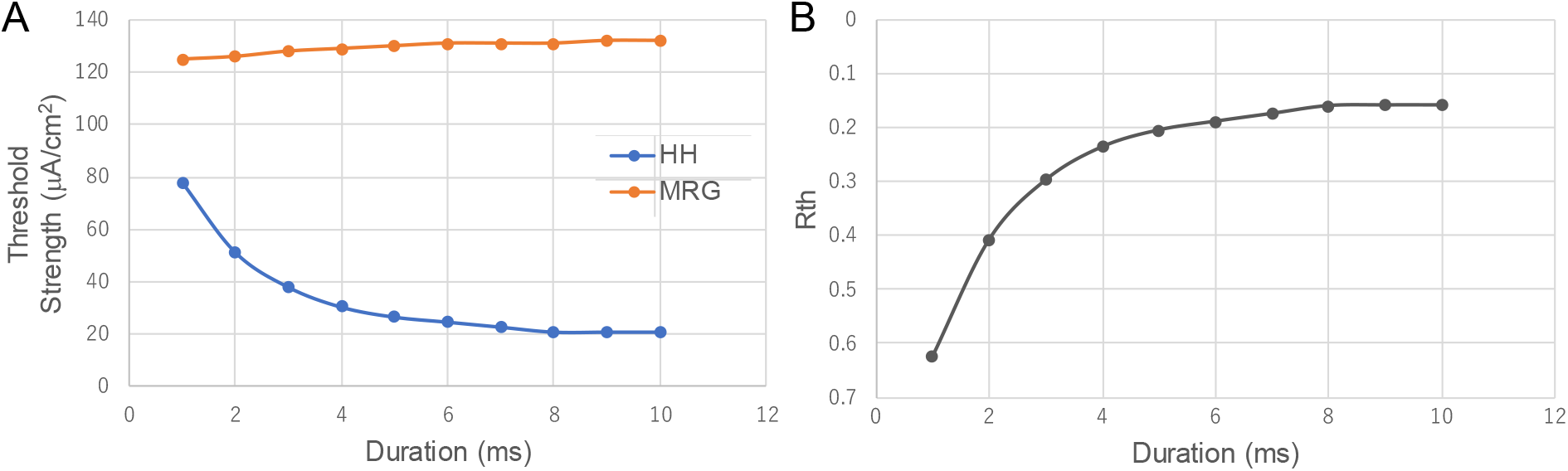
Relationship between threshold strength and anodal duration at the same PAR 1:9. The horizontal axis shows anodal stimulus duration. Note, the total duration of the stimulus is 10 times of the anodal stimulus duration. (A) the threshold strength of *C* and *Aδ*, simulated by the HH and MRG model, respectively. (B) the change of *R_th_*. In this experiment, the ISI between anodal and cathodal stimulus is zero.

#### 3.3.5 Different pulse frequency with different PAR

In addition to duration, we consider that frequency is also a possible factor affecting the selectivity. Two cases were shown in Figure 14. Figure 14A1-C1 show the changes of threshold strength and *R_th_* as frequency increases from 1 Hz to 50 Hz without any limitation to the duration of each period, at PAR 1:6 1:3 and 1:1, respectively. Figure 14A2-C2 show the changes of the same indexes, but with a limitation of 50ms to the maximum duration of each period (i.e., when frequency < 20 Hz, there will be an ISI between each two stimulus periods). For all the PAR values, the threshold strength of the HH model and MRG model didn’t change before 20Hz, while the threshold strength of HH model increased and MRG model decreased after 20Hz in Figure 14A1-C1. As shown in Figure 14A2-C2, even with the limitation to maximal duration of each period, the HH model showed similar changes with those in Figure 14A1-C1, except that, the threshold strength of the MRG model rose at above 15 Hz, and decreased again after 20 Hz.

**Figure 14.**
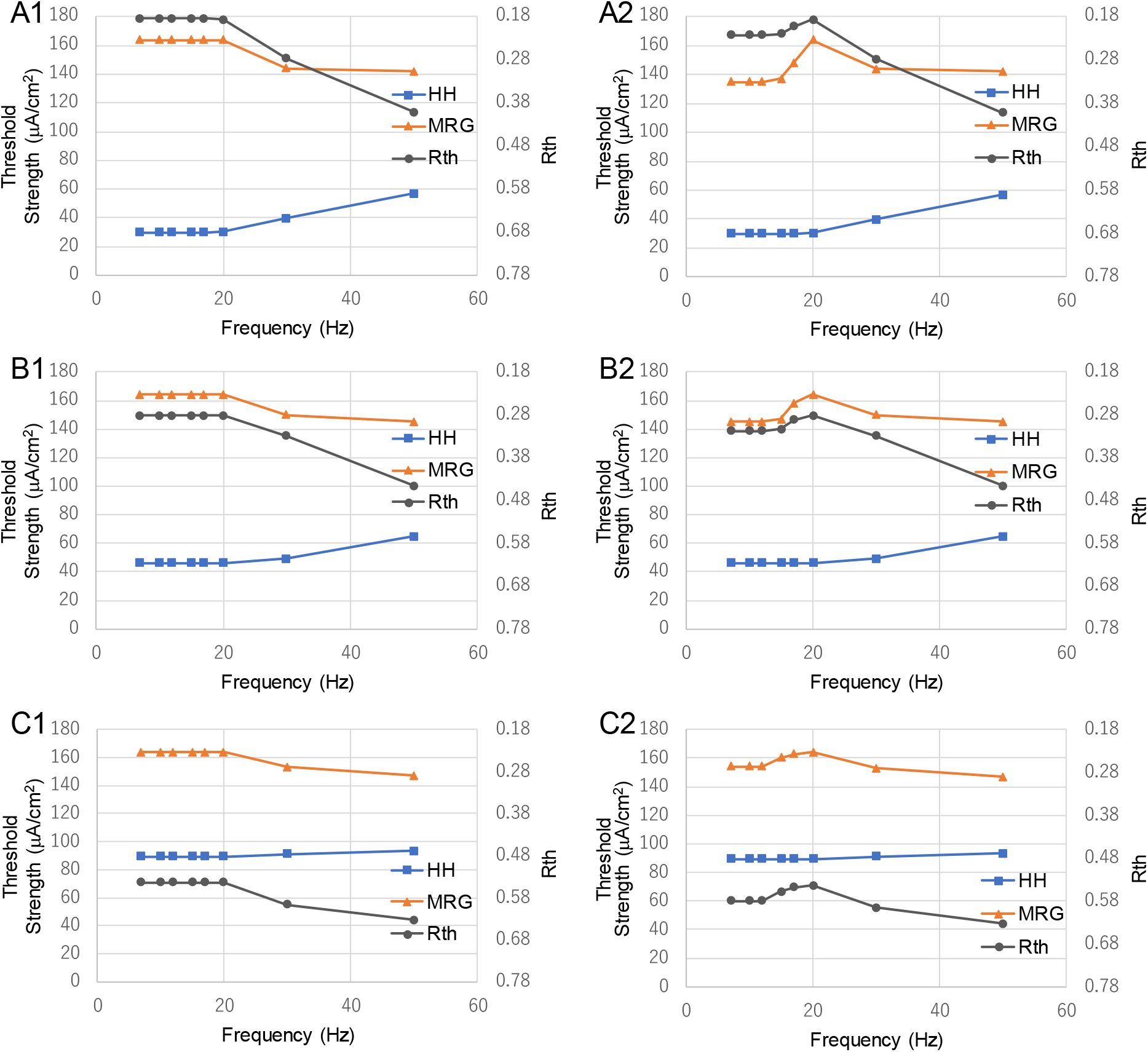
Relationship between pulse frequency, threshold strength and *R_th_* with PAR. (A1) PAR with 1:6, (B1) PAR with 1:3, (C1) PAR with 1:1, (A2) Maximum duration 50 ms, PAR with 1:6, (B2) Maximum duration 50 ms, PAR with 1:3, (C2) Maximum duration 50 ms, PAR with 1:1.

## Discussion

### 4.1 The changes in phase plane caused by preceding anodal stimulation

The threshold potential, its changes and its relative position with respect to the rest point can be visualized clearly in the phase portrait, as illustrated by Figure 3. As seen in the phase portrait of the HH model in Figure 5, when *C* is given an anodal stimulation, the saddle point *b* of the HH model moves downwards closing to the rest point *a* until the two points disappear. During this process, the threshold of *C* gradually decreases and eventually goes below the threshold of *Aδ*. As the duration of anodal stimulation gets longer, *Δd_int* in Figure 5 decreases. When *t_a_* = 6 ms, the intersections disappear, instead the distance between two isocline curves, i.e., *Δd_sep,* starts to increase. The decrease of *Δd_int,* or the increase of *Δd_sep* reduces the activation threshold of *C*, which can be justified by the results of Figure 10A, and Figure 13A, showing the relationship between threshold strength of both nerves and the duration of polarity symmetric and asymmetric stimulation, respectively.

The disappearance of the intersections in phase portraits could be confirmed by comparing different types of waveforms, in terms of the role of preceding anodal stimulation. In all the resultant phase portraits shown in Figure 6, the waveforms with anodal stimulation include C1 (PAR 1:1), D1 (PAR 1:9), and E1 (PAR 9:1), however, only C1 (Figure 6C2: *Δd_HH_sep_* = (0.2, 0.01187)) and D1 (Figure 6D2: *Δd_HH_sep_* = (0.7, 0.02588)) caused the intersection disappearance, whereas, the intersections remain after the beginning of cathodal stimulus (denoted by a red dot) in the phase portrait of E1 ((Figure 6E2: *Δd_HH_int_* = (0.0, 0.00066)), as well as in those of the waveforms without anodal stimulus, i.e., A1 (Figure 6A2: *Δd_HH_int_* = (2.7, 0.01402)), B1, F1. In fact, intersections remain until PAR decreases from 9:1 to 7:1.

The *Δd_sep* in Figure 6D (*t_a_* = 10ms) is larger than that of Figure 6C (*t_a_* = 6ms), and intersections occur in Figure 6B (*t_a_* = 2ms), but its *Δd_int* is smaller than that of Figure 6A (no anodal stimulation case). Referring to the comparison results about duration of preceding anodal stimulation (section 3.3.1, Figure 10), i.e., longer duration causes lower excitation threshold, it is reasonable to state that, the stimulations resulting in the separation of the isocline curves in phase portraits benefit more excitability of *C* than those cause intersected isocline curves, and a larger *Δd_sep* indicates an easier excitation of the nerve.

The importance of the phenomenon could be made clear through crosschecking the ion currents shown in Figure 7, in which the transmembrane ion current right before the excitation of the nerves corresponding to that in Figure 6E1 is almost the same as that of in Figure 6A1, though, quite different from those in Figure 6C1-D1, despite that the case in Figure 6E1 has much longer anodal duration than that of Figure 6C1-D1 under the same charge constraint. The stimulation with PAR 7:1, which is the border value to cause disappearance of intersections, has a comparatively large difference from that of PAR 9:1 (the difference of *ΔI*: 5.99 μA, larger than the maximal *ΔI_Na_*: 4.49 μA, caused by any other PAR values). Thus, it is reasonable to state that the disappearance of intersections *a* and *b* did play a role in the reduction of potassium ion current, hence the reduction of activation threshold of the *C*.

Moreover, a larger *Δd_HH_sep_* in D2 (0.7, 0.02588) compared with that of C2 (0.2, 0.01187), is caused by its stronger anodal stimulus in D1 (180 μA/cm^2^) compared with that in C1 (86 μA/cm^2^), as the duration of the anodal stimulus of C1 and D1 is same (10 ms). This was further confirmed by the fact that E1 has lower intensity than both C1 and D1, thus could not cause the disappearance of intersections in the phase plane. In other words, as the intensity of anodal stimulus increases, the distance *Δd_HH_sep_* between the two isocline curves becomes larger, which indicates a stronger effect on a reduction in activation threshold.

Furthermore, since the polarity asymmetric waveforms do not affect the activation threshold of *Aδ* as much as that of the *C*, as reflected by the comparison between their behavior in phase plane that is shown in Figure 6A3-F3 and Figure 6A2-F2.

### 4.2 Ion channel variable and current analysis

Specifically, the threshold strength, i.e., the intensity of the external stimulus necessary for eliciting action potential, is affected by the state of each ion channel before the membrane potential exceeds its threshold. It can be seen from equation (1) that the reduction of external cathodal stimulation intensity means that a higher current from the ion channel is required to compensate for generating the action potential. Thus, the three ion channel gate variables are the focus of analysis. In Figure 8A which shows the phase portrait of membrane potential *v* and sodium channel gate variable *m,* no matter where at the end of the anodal stimulation the initial value of *m* is located at, the trajectories from the beginning of the cathodal stimulation (denoted by red dots) to the generation of the action potential(denoted by black dots) are all on the almost same curve for different stimulation waveforms. Therefore, anodal stimulation does not affect fundamentally the behavior of the variable *m*. This is because the small time constant of *m* of both HH and MRG model, as shown in Figure 2, leads to their fast-changing sodium channel, it is difficult for the difference caused by anodal stimulation to affect the rising of *m* during cathodal stimulation.

Compared with the variable *m*, the changes of the variables *h* and *n* are very significant. In the phase portraits of *v-h*, and *v-n*, different stimulation waveforms (cathodal stimulation only, PAR 1:1-1:9) resulted in clearly different trajectories. It can be seen from equation (2) that a higher *h* increases the current of the sodium (Na) ion channel, and a smaller *n* reduces the current of the potassium (K) ion channel, which increases the current flowing from the outside to the inside of the membrane, and reduces the current flowing outward the membrane, respectively, so that the action potential can be achieved with lower cathodal stimulation.

As stimulus changes from cathodal-stimulation-only to bipolar PAR 1:1, 1:3, 1:6, further to 1:9, the gap between the phase portraits of each two neighboring stimulation waveforms gradually decreases, which agrees with the decreasing trend of the threshold of the HH model in Figure 12A. That is, although a smaller PAR results in a bigger reduction of the cathodal stimulation threshold, further decreasing the PAR, may cause saturation of the improvement in *C*-selectivity.

Figure 9 shows the changes of the sodium (Na) ion channel current and the potassium (K) ion channel current before the threshold potential. Compared with the significant gap of variable *h* in Figure 8B, the gaps between the Na ion currents at threshold potential corresponding to different stimulation waveforms in Figure 9A is not very obvious (average: 10.21573 μA, standard deviation: 1.14962 μA). In contrast, the gaps between K ion currents in Figure 9B is very significant (average: 72.56375 μA, standard deviation: 9.25445 μA). It can be seen from equation (2) that the Na ion current is positively correlated with the cube of *m* and *h*. Moreover, because before the threshold potential, the value of the variable *m* is very small, the cube of *m* becomes an even smaller value, which greatly reduces the gap between different variables *h* (Figure 8A). On the other hand, the K ion current is positively correlated with the fourth power of *n*, which further enlarges the gap caused by the difference in the variable *n*. Therefore, it is the K ion channel (regulated by the variable *n*) that plays a decisive role in the reduction of the activation threshold of *C*.

One issue to note is that the K ion currents (*I_K_*) of different waveforms show different behaviors after and before the membrane potential exceeds the threshold potential. As shown in Figure 9C, after its membrane potential reaches the threshold potential, *I_K_* corresponding to the waveform PAR 1:3 is much higher than that of the others, whereas, all the K ion currents did not show a difference, before threshold potential. Since the time to reach the action potential from its rest state only changed slightly with threshold strength, *I_K_* does not affect the process after the model reaches the threshold potential, even though it does affect the speed from the threshold potential to the action potential.

In summary, the introduction of anodal stimulation can effectively lower the threshold of the unmyelinated nerve, while it does not affect so much the threshold of the *Aδ*. Although it has been suggested that it is the anode break in peripheral sensory nerves that may cause double peak potentials [PedroPereira 2016, Therimadasamy 2015, 39 40], which had been previously thought to be elicited by depolarization of nerve terminal axons or skin receptors [Aprile 2007, 41], the underlying ion mechanism has not been understood yet. Moreover, the double peak potentials were usually generated by stimulation with low intensity, long duration, and the latency of the second peak was much longer than the first peak generated by cathodal stimulation [Aprile 2003, 42]. These stimulation conditions are consistent with the favorable stimulation conditions for *C* in our research. Our study showed that if the double peak potentials are caused by anode break, then it is the anodal stimulation to *C* that generated them.

And a lower PAR can also widen the threshold strength gap. It can be declared that the use of anodal polarity asymmetric stimulation has a positive impact on selective stimulation of nociceptive nerve fibers. However, as shown in Figure 7 and Figure 12, as the PAR is getting lower, the effect of polarity asymmetric stimulation on the reduction of the threshold strength of the unmyelinated nerves turns to saturate. And, for a charge balanced stimulation, a lower PAR requires higher intensity of anodal stimulation, which might excite surrounding tissues such as muscle and *Aβ* nerve fibers that are related to sensation of touch, pressure, and vibration.

### 4.3 The effect of stimulation waveform parameters on *C*-selectivity

Polar precedence, duration and ISI and polarity asymmetry related to the bipolar stimulation were investigated in detail. In the experiment on the polar precedence, the results about the cathodal-first stimulation (Figure 10B) shows that, the threshold strength of the *C* is reduced, and closed to the threshold strength of the *Aδ*, but cannot be lower than it, which is in agreement with those from the other simulation studies and animal experiments [Gaines Jessica L. 2018, Boyd IA 1979, 31 43], in which the anodal stimulus was given after the cathodal stimulus to balance the charge-injection for safety consideration. The results here were consistent with the general understanding that thicker fibers are more likely to be stimulated to produce action potential than the thinner ones [Grill WM 1997, 15]. The threshold strength ratio of the responses to stimulation agrees well with the time constants of ion channel variables of the *C* and *Aδ* that are shown in Figure 2. As shown in Figure 2, the time constants of *m* and *h* in HH model (the highest value 0.5014, 8.5820, respectively, in Figure 2A) are higher than those of *m* and *h* in MRG model (the highest value 0.2167, 1.1510, respectively, in Figure 2B), which means that HH model needs much more time to adapt to external stimuli.

In contrast, anodal-first stimulation favors *C*-selectivity, and pushed the threshold of the *C* to even below than that of the *Aδ*. This phenomenon is similar to anode break [Ranjan 1998, 21], which showed the possibility of eliciting action potentials by only anodal stimulation. However, neither the role of the charge balancing following cathodal stimulation, nor the promotion of *C* excitation and *C*-selectivity over *Aδ* have been addressed. Through the phase portrait analysis shown in Figure 5, it is clear that the preceding anodal stimulation with duration long enough, in conjunction with a following cathodal stimulation, can have a significant effect of promoting *C*-selectivity.

Moreover, though, the duration of anodal stimuli strongly affects the promotion effect of *C*, the threshold strength of *C* is not linearly dependent on the duration. Compare the results shown in Figure 10A and Figure 13A, a low PAR could greatly reduce the threshold strength of the HH model (PAR: 1:1-1:9, threshold strength: 87.4-20.9 μA/cm^2^), but they need the same duration (anodal duration: 8 ms) of stimulus to reach the minimum threshold strength. For the same total duration of stimulus such as 10 ms, the PAR 1:1 (anodal duration:5 ms, threshold strength: 95.0 μA/cm^2^) and a much lower PAR, PAR 1:9 (anodal duration:1 ms, threshold strength: 77.9 μA/cm^2^), did not show a large difference (17.1 μA/cm^2^) in threshold strength. However, as the total duration turns longer such as 30 ms, the case of PAR 1:1 (anodal duration:15ms, threshold strength: 87.4 μA/cm^2^, which is the same with all the cases with duration longer than 8ms, according to the HH threshold strength results shown in Figure 10A) and the one with a lower PAR, PAR 1:9 (anodal duration:3ms, threshold strength: 38.0 μA/cm^2^) showed a large difference (49.4 μA/cm^2^). Thus, only when the anodal stimulation duration is long enough, the PAR can significantly improve *C*-selectivity.

On the other hand, neither anodal-first nor cathodal-first stimulation significantly promoted the *Aδ*. When the stimulation duration increases, the charge of the anodal stimulation also increases. That is, it needs more charge of the cathodal stimulation to counteract the anodal stimulation effect for the *Aδ*, thus, the threshold strength of the *Aδ* slightly increases. Although the effect of charge accumulation also affected *C*, it is offset by the promotion effect of anodal stimulation on *C*. In addition, the change of *R_th_* shown in Figure 13B gradually decreases, which implies that the stimulation with extremely long anodal duration (i.e., > 8 ms) can hardly contribute to a further reduction of *R_th_*. Notice for PAR 1:1 in Figure 10, the *R_th_* also saturated at 8 ms anodal duration, it is reasonable to have duration of anodal stimulation shorter than 8ms.

The PAR of stimulation is related to the preceding anodal stimulation in terms of both intensity and duration, thus its effect on the threshold strength of *C* is nonlinear, too. There have been studies reporting the effects of pre-pulses with different intensity and duration on neuro dynamics [Bostock 1998, 19]. However, in the literature, PAR has not been studied as a comprehensive parameter on its effect on *C*-selectivity. In one relevant study [Cogan SF 2006, 44], the effect of charge injection with asymmetrical waveforms was investigated through in vitro experiments. Though, neither its effect on the excitability of C, nor the mechanism behind has been addressed. Thus, the PAR, with its role and underlying mechanism identified in this study, can be a new dimension for designing effective selective stimulation.

Responding to the stimulation with increasing anodal duration, and decreasing intensity (a higher PAR value), the threshold strength of *C* gradually rises and exceeds that of *Aδ*, while the threshold strength of *Aδ* decreases slightly and remains unchanged after 5:1, as shown in Figure 12B. It can be seen from Figure 12C that *R_th_* changes rapidly around PAR 1:1. This is also consistent with the change of ion current or potassium channel parameter *n* in Figure 7A and Figure 8C. Since the duration of the cathodal stimulus is constant in the experiments shown in Figure 12A, an increase in the threshold strength of the *C* also indicates that the charge (or energy) accumulated by the cathodal stimulus is increased (Figure 12D). In contrast, the threshold strength of *Aδ* decreases as the pulse width of anodal stimulus increases, and its intensity decreases. As shown in Figure 12A, since the duration of cathodal stimulus of all waveforms are same, it was the anodal stimulus that made the difference. Furthermore, because the effect of duration saturated at a certain value as shown in Figure 10, the intensity of the anodal stimulus is playing the major role. However, a stronger anodal stimulation may have potential safety concerns. Regardless of anodal and cathodal stimuli, excessive stimulation intensity may affect other subcutaneous tissues or even cause damage to them [Mallik A, 2005, 45]. From the anodal duration-*R_th_* graphs for PAR 1:1 (Figure 10C) and PAR 1:9 cases (Figure 13B), the improvement of *R_th_* might be saturated at anodal duration of 8 ms. The anodal stimulus longer than 8 ms can have a better stimulation effect of *C*, and a lower asymmetric ratio can lead to better *C*-selectivity (Figure 12C). However, when the above two conditions are met simultaneously, the stimulation waveform may be too long, which might cause safety issues [Merrill DR 2005, 46]. Therefore, even though waveforms and parameters for achieving better *C*-selectivity were identified in this study, the range of the key parameters, such as the intensity and pulse width of the anodal stimulation, needs to be further investigated to ensure the safety and effectiveness of *C*-selectivity for surface stimulations.

As shown in Figure 14, the relationships between threshold strength, *R_th_* and pulse frequency at three different PAR values are consistent with those shown in Figure 12. For the three PAR 1:6, 1:3, 1:1, the value of *R_th_* mostly depends on the threshold strength of the HH model. In Figure 14A1-C1, since there is no restriction on the stimulation duration, the threshold of the MRG model remains the same as HH model before 20 Hz. On the one hand, long-term stimulation will cause the Faradaic charge transfer which leads to safety hazards [Merrill DR 2005, 46], on the other hand, the stimulation frequency less than 20Hz does not further improve the *C*-selectivity, we limited the stimulation duration to 50 ms in Figure 14A2-C2. The result shows that the threshold of the HH model is unchanged, but the threshold of the MRG model is reduced at low frequencies and returned their maximum at 20 Hz, regardless of the PAR value. For the *R_th_* of each PAR, the main changes came from the increase of threshold strength of the MRG model when stimulated at frequencies larger than 15 Hz, 15 Hz and 12 Hz, for PAR 1:6, 1:3, and 1:1, respectively. A shorter interstimulus interval resulted from that frequency band has a stronger effect on increasing the threshold strength of the MRG model. This may be due to the extremely large time constant of parameter *s* in the MRG model [McIntyre 2002, 23]. However, when the frequency is greater than 20 Hz, the threshold strength of the MRG model begins to decrease, while the threshold strength of the HH model increases. Especially at a low PAR (1:6), the magnitude of the change is the largest. As the frequency further increases, the *R_th_* of Figure 14A2 (PAR 1:6) changes more than that in Figure 14B2-C2, making smaller difference in their *R_th_*. This reflects that when the frequency is getting close to the higher frequency band (>20 Hz), the influence of frequency is larger, otherwise it becomes smaller. Although other clinical or medical research also used low-frequency electrical stimulation [Harris, G. W. 1969, 10], after comparing Figure 14A2-C2, lower frequency will amplify PAR’s effects on *R_th_* (the difference of *R_th_*: 0.09616 between PAR 1:6 and 1:3 with 20 Hz, is larger than the difference of *R_th_*: 0.04411 between PAR 1:6 and 1:3 with 50 Hz). Therefore, when using stimulation with long total (anodal and cathodal stimulation) duration (50-100 ms from Figure 13A), selecting a low PAR and higher frequency (15 Hz-20 Hz) in the lower frequency band can improve the *C*-selectivity. It is necessary to trade off the *C*-selectivity effect and safety when performing stimulation below 15 Hz. Though, as reported by the other studies [Siyu He, 2020, 14], in the higher frequency band (100-500 Hz), however, the frequency turns to be an important factor to promote the excitation of *Aδ* while inhibiting *C*.

Different from the interval between two bipolar stimuli, the ISI between an anodal pulse and a cathodal pulse has an impact on the safety of stimulation. It has been shown that this ISI needs to be less than 10 ms and greater than 6ms to meet the safety requirements of bipolar stimulation [Mohamed MA 2017, 47]. As stated in [Mohamed MA 2017, 47], the aim of adding ISI into the cathodal-first stimulation waveform is to reduce threshold strength of the following anodal stimulation. However, in order to guarantee the safety of stimulation, the interval between cathodal and anodal stimuli cannot be too large. Therefore, choosing an appropriate duration of ISI is important to improve the effect of the cathodal-first stimulation. On the other hand, for the bipolar anodal-first stimulation, according to our results shown in Figure 11, a shorter ISI or even no ISI can maximize the selective stimulation effect. A shorter ISI can not only improve the *C*-selectivity, but also reduce the impact on the organs. This can be understood from the perspective of safety as follows. The anodal stimulation and cathodal stimulation should be as close as possible to ensure that the charge can be neutralized within a short time.

In addition, the interval between each bipolar pulse has an effect on the stimulation results (such as the effect of refractory period reported in [Dudel J. 1983, 48]). Obviously, for pulses with a low frequency of 1 Hz, nerves stimulated by the current bipolar pulse are less subjected to the influence of its previous bipolar pulse, since there is enough time for the neuron to recover (Figure 14A2-C2).

### 4.4 Contribution and limitations

In this study, prepulse-based approach and frequency-based approach were explored and integrated for the purpose of realization of high *C*-selectivity over *Aδ*. The contribution of this paper can be summarized as follows.

1. Through investigating the responses of the simulation models of two types of nociceptive nerve fibers in a phase plane and their ion currents, the effectiveness of the anodal-first stimulation for *C*-selectivity, and its ion channel mechanism were made clear for the first time in this research area. Bipolar square waves are not just for charge balancing, with appropriate intensity and duration of anodal stimuli, it could reduce significantly the potassium current flowing out through the membrane, thus lower the threshold strength of *C*. Moreover, by using the index for *C*-selectivity, *R_th_*, and index in the phase plane, *Δd_int* and *Δd_sep,* it turned to be clear that, for the polarity asymmetric pulses, intensity of anodal stimulus is a more important factor than its duration, though, a certain duration of the anodal stimulus is necessary to guarantee the *C*-selectivity.
2. The effects of the important parameters for continuous periodic bipolar stimulation, on *C*-selectivity over *Aδ* were first investigated. The parameters include frequency, duration, and ISI, and polarity asymmetry ratio (PAR). The landscape of the solution space of *C* selective stimulation was made clear. For the polarity symmetric stimulation, 20 Hz is the best frequency for *C*-selectivity. Lower PARs are better for *C*-selectivity in terms of both *R_th_* and potassium ion currents. However, considering the safety of stimulation, there might be a constraint on the intensity of the anodal stimuli.

Besides contributions, limitations were noticed as well. Since the ultimate goal is to realize surface selective stimulation, the influence of skin should be taken into consideration. It is beyond the scope of this paper, though, definitely, this needs to be made clear in near future. Moreover, the influence of the polarity asymmetric bipolar stimulation on other nerves and tissues except *C* and *Aδ* was not clear, which is required to be considered and reflected in the simulation models. Furthermore, the findings from the simulation study need to be carefully verified, and further validated. A close cooperation with both subjective questionnaire and objective evaluation, such as studies on pain-related electrical-stimulation evoked electroencephalogram [Tripanpitak K. 2020, 49] is crucial.

## Conclusion

In order to make clear how a preceding anodal stimulation and a cathodal stimulation (named as anodal-first stimulation) could lower the activation threshold of nociceptive *C* over myelinated nociceptive *Aδ*, and further clarify the landscape of the solution space, the HH model for *C* and the MRG model for *Aδ*, were employed for comparing their responses to relevant waveforms in terms of both their behavior in phase plane, and the *C*-selectivity. It was made clear that the anodal-first polarity asymmetric stimulations are more likely to stimulate the unmyelinated nerves. This is because that the preceding anodal stimulation could decrease the potassium ion current of them for the following cathodal stimulation. The optimal parameters in terms of activation threshold have been identified in the low frequency band, which showed especially high possibility of *C*-selectivity. This is an important step towards the long-term pain relief for chronic pain.

## Acknowledgements

Siyu He was supported by the State Scholarship Fund awarded by the China Scholarship Council (CSC) (Grant number: 201708050123). We also gratefully acknowledge other members of our laboratory for their kind help.

## Competing interests

We claim that this work is partially supported by a joint-research project between Chiba University and Omron Healthcare Co., Ltd., and Shozo Takamatsu is an employee of Omron Healthcare Co., Ltd.

## Appendix I

HH model parameters:

|  |  |
| --- | --- |
| Maximum $Na^+$ channel conductance ($g_{Na}$) | 120.0 mS/cm<sup>2</sup> |
| Maximum $K^+$ channel conductance ($g_K$) | 36.0 mS/cm<sup>2</sup> |
| Leak channel conductance ($g_{Lk}$) | 0.3 mS/cm<sup>2</sup> |
| Membrane capacitance (C) | 1.0 uF/cm<sup>2</sup> |
| $Na^+$ reversal potentials ($E_{Na}$) | 50.0 mV |
| $K^+$ reversal potentials ($E_K$) | -77.0 mV |
| Leak reversal potentials ($E_{Lk}$) | -54.4 mV |
| Rest potentials | -65.0 mV |

Parameters from equation (3) for HH model:

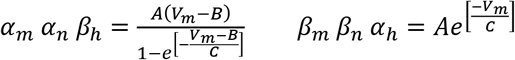

| Voltage and time dependent parameters | A(m/s) | B(mV) | C(mV) |
| --- | --- | --- | --- |
| $\alpha_m$ | 0.10 | 25.00 | 10.00 |
| $\beta_m$ | 4.00 | 0.00 | 18.00 |
| $\alpha_h$ | 0.07 | 0.00 | 20.00 |
| $\beta_h$ | 1.00 | 30.00 | 10.00 |
| $\alpha_n$ | 0.01 | 10.00 | 10.00 |
| $\beta_n$ | 0.125 | 0.00 | 80.00 |

## Appendix II

MRG model parameters:

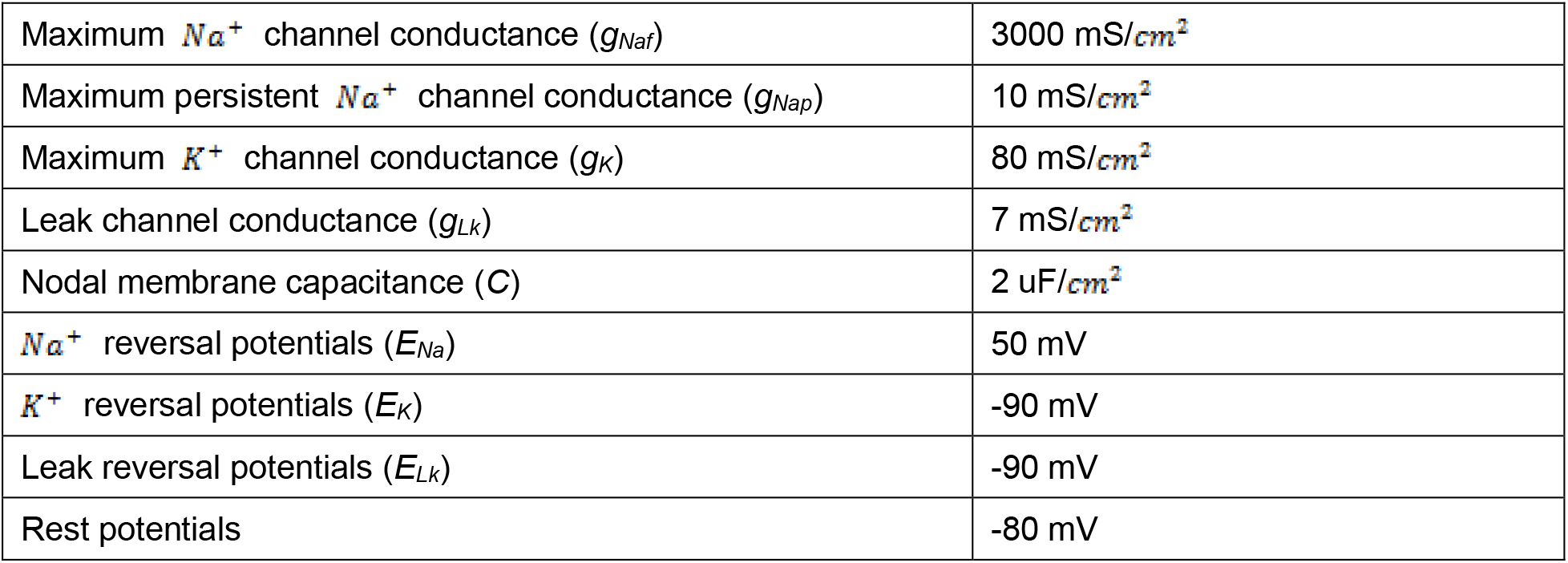

Parameters from equation (3) for HH model:

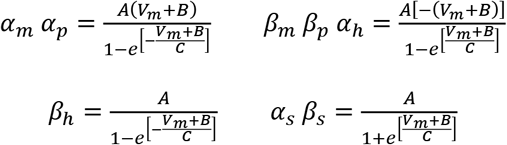

| Voltage and time<br>dependent parameters | A(m/s) | B(mV) | C(mV) |
| --- | --- | --- | --- |
| $\alpha_m$ | 6.57 | 20.40 | 10.30 |
| $\beta_m$ | 0.304 | 25.70 | 9.16 |
| $\alpha_p$ | 0.0353 | 27.00 | 10.20 |
| $\beta_p$ | 0.000883 | 34.00 | 10.00 |
| $\alpha_h$ | 0.34 | 114.00 | 11.00 |
| $\beta_h$ | 12.60 | 31.80 | 13.40 |
| $\alpha_s$ | 0.30 | 53.00 | -5.00 |
| $\beta_s$ | 0.03 | 90.00 | -1.00 |

